# Characterizing Sequence-Function Relationships in Chimeric DcuS/EnvZ Histidine Kinases at Scale

**DOI:** 10.64898/2026.06.03.728855

**Authors:** Luca B. Lippert, Samuel R. Hinton, Andrew S. Holston, Karl J. Romanowicz, Calin Plesa

## Abstract

While bacterial sensor histidine kinases (SHKs) are widespread as natural molecular biosensors, tools for high-throughput characterization of SHK signaling phenotypes are limited, hindering wide scale implementation of bacterial-based sensing. Here, we developed a synthetic two-component signaling system that reports chimeric SHK signaling via a standardized fluorescence readout. With this synthetic system, we screened a library of chimeric DcuS/EnvZ SHKs to characterize sequence-function relationships within in the DcuS sensory and transmembrane domains. We quantified the effects of 1,173 mutations on signaling outputs in the presence of fumarate, a native DcuS ligand, as well as aspartate for which DcuS has minimal affinity for. We identified eleven positions across the DcuS domains which significantly alter aspartate responsiveness and selectivity and further observed a role for cytoplasmic N-terminal residues in determining signaling outputs. In future studies, this framework will expedite design of biosensors for novel ligands by enabling high-throughput screening of mutagenized libraries of natural SHKs.

## Introduction

Biosensors are foundational tools in synthetic biology, enabling environmental sensing,^1^ dynamic pathway regulation,^2^ theragnostics,^3^ metabolic flux analysis,^4^ and high-throughput screening applications.^5^ Expanding the repertoire of genetically encoded sensors and engineering ligand specificity in existing receptors remain major goals in the field. One mechanism through which organisms detect and respond to changes in their environments is via two component systems (TCS), which are used by prokaryotes^6^ and eukaryotes^7^ alike. TCSs consist of a sensor histidine kinase (SHK) and cognate response regulator (RR).^8^ SHKs and RRs are two of the largest paralogous gene families found in bacteria, with many species encoding for dozens of TCSs, such as *Bacillus subtilis* (*B. subtilis*),^9^ *Escherichia coli* (*E. coli*),^10^ *Pseudomonas aeruginosa*,^11^ and cyanobacteria.^12^ Strikingly, the multicellular bacterium *Myxococcus xanthus* encodes for over 260 TCS proteins,^13^ highlighting the potential sensory repertoire possible in an engineered chassis.

The general mechanism of signal transduction in TCSs is phosphotransfer from the SHK to the RR upon stimulus detection.^14^ With this core signaling logic, SHKs and RRs are capable of integrating a wide range of external signals to determine cellular behavior. SHK proteins are typically homodimeric and utilize a highly variable extra-cytoplasmic sensory domain for detecting diverse stimuli, including simple sugars,^15^ amino acids,^16^ nitrate,^17^ quorum sensing autoinducers,^18–20^ antimicrobial peptides,^21^ osmolality,^22^ temperature,^23^ and specific wavelengths of light.^24^ Upon stimulation, the SHK transduces the signal through its transmembrane (TM) domains to the cytosolic CA (<u>c</u>atalytic and <u>A</u>TP binding) and DHp (<u>d</u>imerization and <u>h</u>istidine <u>p</u>hosphotransfer) domains of the SHK.^14^ In canonical TCSs, signal perception upregulates autophosphorylation onto a conserved histidine residue in the DHp domain and subsequent phosphotransfer to its cognate RR,^8,25,26^ with coevolving residues in both proteins conferring specificity to the interaction.^27,28^ The phosphorylated RR then alters gene expression or catalyzes the modification of other proteins^29^ to drive cellular behaviors such as chemotaxis,^30^ nutrient uptake,^31^ cell division,^32^ stress responses,^33^ virulence,^34^ and more. Furthermore, most SHKs are possess bifunctionality as both a kinase and phosphatase, and ligand-dependent signaling outputs derive from changes to the net rate of RR phosphorylation.^22,35–38^

With great functional diversity comes great sequence diversity: approximately 18 million SHK sequences have been putatively identified from bacterial genomes,^40^ but comparatively few TCSs have been thoroughly characterized. This imbalance represents a major bottleneck in our ability to understand, predict, and engineer bacterial signaling. In particular, we still lack scalable approaches for determining how residues across TM and sensory domains shape downstream signaling outputs. Establishing tools to characterize sequence-function relationships in currently known SHKs at scale would enable the development of a diverse array of new biosensors with tunable signaling phenotypes, allowing synthetic biology to harness this large repository of bacterial receptors towards novel applications.

Native TCSs operate within complex regulatory networks, making it difficult to dissect sensory inputs and outputs from downstream signaling contexts;^5^ however, the modular separation of input detection, phosphotransfer, and output regulation between domains in TCSs provides a structural basis for functional reprogramming.^41^ Previous work dating back to 1989 has demonstrated the utility of SHK and RR chimera engineering for synthetic biosensor design (for review, see ref. 42).^42–47^ This approach can be scaled to systematically discover and engineer SHKs for novel signals by rewiring variable SHK sensory and TM domain signaling to a standardized, measurable output.^48^ EnvZ is among the most well studied SHKs and has been used as a kinase fusion partner in many chimeras, making it a strong choice for a modular chimera engineering platform. Such a system would be capable of associating SHK sequences with inputs with modern high-throughput techniques, expediting the identification of functional determinants at residue-level resolution.

Building on this framework, we establish scalable methodology for chimeric SHK characterization, and apply this system to tune the signaling phenotype of the C4-dicarboxylate sensor, DcuS,^49^ by screening a mutagenic library of DcuS sensory and TM domains fused with the CA and DHp domains of EnvZ. To generate a standardized output for DcuS/EnvZ chimeras, we built on an existing synthetic RR reporter system, wherein a chimeric RR, OmpR/CcaR, modulates expression of superfolder GFP (sfGFP),^50^ to report DcuS/EnvZ activation. We confirmed that the DcuS/EnvZ chimera is strongly activated by fumarate, the endogenous DcuS ligand, but not aspartate, for which native DcuS shows low affinity. We next leveraged the synthetic TCS to characterize the mutational landscape of the DcuS sensory and TM domains in this chimeric context. We identified eleven positions in DcuS that significantly increase aspartate specificity and reshape overall signaling output, and further validated this approach by characterizing individual mutants. More broadly, this work establishes a scalable platform for mapping SHK sensory and TM domain sequence to signaling phenotype, providing a path toward large scale SHK characterization and the engineering of bacterial biosensors with tunable input-output behaviors.

## Results and Discussion

### Design of synthetic TCS

In order to develop a scalable methodology for characterizing new biosensors, we first had to clone a functional DcuS/EnvZ chimera and, secondly, quantify its activation via a synthetic TCS. Previous work^46^ established a functional DcuS/EnvZ chimera composed of the TM1, sensory, and TM2 domains of DcuS and the CA and DHp domains of EnvZ, and was constructed here by fusing coding sequences for DcuS^1–193^ with EnvZ^231–450^ (Fig. 1E). To determine if DcuS/EnvZ chimera activity can be reported via fluorescence, the cloned DcuS/EnvZ plasmid was co-transformed with a reporter plasmid, pSR40.29, which encodes for OmpR/CcaR and an OmpR/CcaR inducible sfGFP gene (Fig. 1A). Importantly, the DNA-binding domain of OmpR/CcaR was mined from the genome of *B. subtilis* and is orthogonal to the native *E. coli* machinery,^50^ allowing for isolation of the synthetic TCS. Additionally, the synthetic TCS was implemented in a derivative of *E. coli* strain BW29655 (WT), a Keio collection strain, that lacks genomic copies of *envZ* and *ompR,*^51^ selected to further reduce crosstalk with native *E. coli* systems. This resulted in strains BW29655-pSR40.29 (pSR) and BW29655-pSR40.29-pDcuSEnvZ (synTCS) (Table 1).

**Figure 1.**
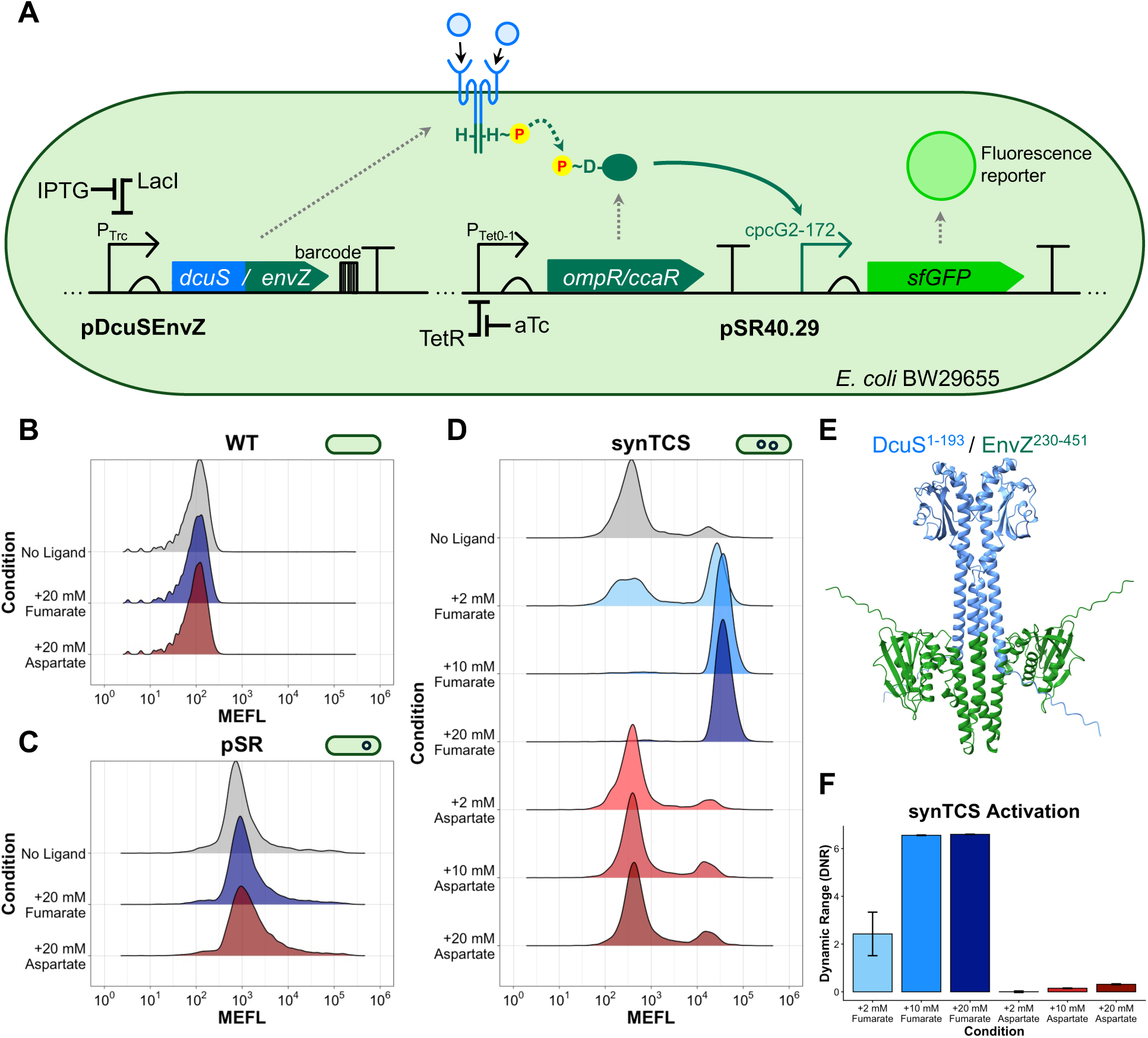
The DcuS/EnvZ chimera and synthetic TCS system. (A) Schematic overview of synthetic TCS and fluorescence reporter. (B) Flow cytometry analysis of WT singlets grown without ligand, 20 mM fumarate, and 20 mM aspartate. (C) Flow cytometry analysis of pSR singlets grown without ligand, 20 mM fumarate, and 20 mM aspartate. (D) Flow cytometry analysis of synTCS singlets grown without ligand or with 2, 10 or 20 mM fumarate or aspartate. (E) AlphaFold2 Multimer structure prediction of the DcuS/EnvZ homodimer. (F) log_2_ of the ratio of ligand to no ligand MEFL fluorescence, or dynamic range (DNR) of synTCS for varying concentrations of fumarate and aspartate.

**Table 1.**
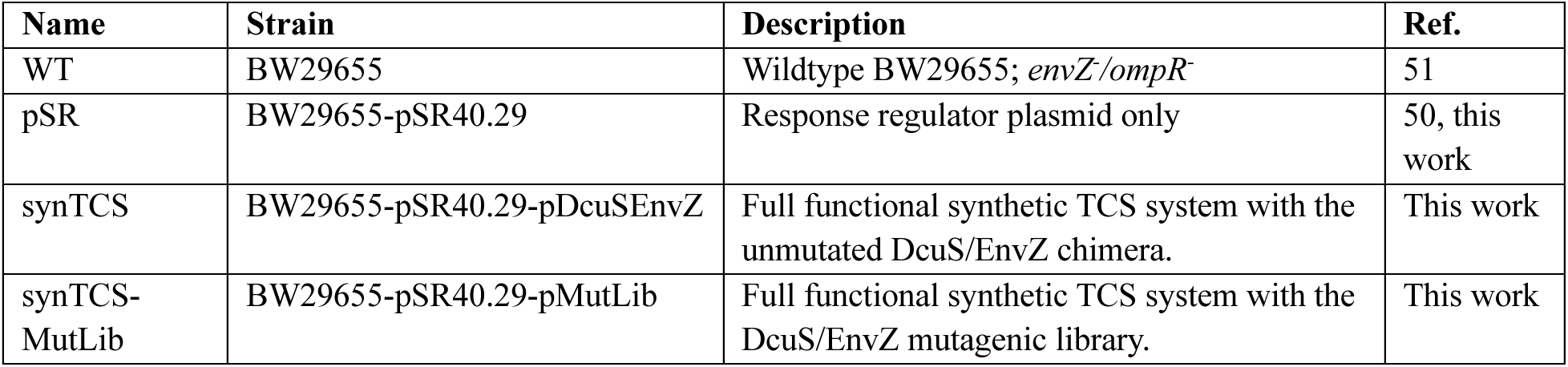
List of BW29655 transformants used in this work.

### Ligand-dependent activation of synTCS

Flow cytometry analysis revealed low autofluorescence of WT cells across all conditions (Fig. 1B, Fig. S1A), while synTCS was strongly activated by fumarate but not aspartate (Fig. 1D, Fig. S1C). Interestingly, synTCS exhibited analog responsiveness to both ligands: dynamic range (DNR; Eq 1, Methods) increased strongly at 10 and 20 mM of fumarate and minorly at the same concentrations of aspartate (Fig. 1F). Aspartate responses by DcuS have been sparsely reported in the literature^15^ and it is generally accepted that DcuS is a nonspecific C4-dicarboxylate receptor.^52^ Our data demonstrates that DcuS is preferentially activated by fumarate and has low affinity for aspartate, confirming that the DcuS/EnvZ chimera retains the native ligand specificity of DcuS.

### DcuS/EnvZ suppresses noncognate activation of OmpR/CcaR

We further observed intermediate fluorescence for single cells of pSR, containing only OmpR/CcaR and sfGFP, irrespective of ligand presence or concentration (Fig. 1C, Fig. S1B). This result led us to investigate how the individual components of the full system in synTCS influence fluorescence readout. First, we observed fluorescence did not increase over time when OmpR/CcaR was not induced, in either pSR (Fig. S1A) or synTCS (Fig. S1B-C). sfGFP expression was increased when OmpR/CcaR but not DcuS/EnvZ was induced in synTCS (Fig. S1D), and induction of DcuS/EnvZ abolished this background fluorescence in the absence of fumarate (Fig. 1C, Fig. S1C).

It was previously observed that native OmpR can be phosphorylated by CpxA^53^ and by small molecule phosphodonors such as acetyl phosphate,^54–56^ providing a plausible explanation for sfGFP expression in the absence of DcuS/EnvZ. The apparent suppression of activated P∼OmpR/CcaR by DcuS/EnvZ without fumarate is consistent with early *in vitro* studies of mutant Tar/EnvZ (Taz) chimeras,^38^ which demonstrated dephosphorylation of P∼OmpR by EnvZ chimeras. Others have shown a high dissociation constant (K_d_) between EnvZ and P∼OmpR,^56^ which may impact dephosphorylation *in vivo*. However, in this case, overexpression from plasmid-encoded constructs may compensate and allow dephosphorylation to occur. Taken together, our data supports a model wherein OmpR/CcaR is noncognately phosphorylated when DcuS/EnvZ is not expressed, resulting in a fluorescence signal; upon induction, DcuS/EnvZ sequesters and dephosphorylates P∼OmpR/CcaR, restoring ligand-dependent control of reporter expression.

### Flow-seq enables genotype-phenotype associations in synTCS-MutLib

With our driving motivation being the development of chimeric SHKs with novel signaling phenotypes, we next applied the synthetic TCS to identify mutations that alter the signaling output of DcuS by creating a mutant DcuS library and evaluating the mutants in a high-throughput screen. To create the mutant library, error prone PCR was performed on the DcuS TM1, sensory, and TM2 domains, and the pool of mutagenized DcuS sequences was cloned and transformed into pSR for a final library diversity of 1.5×10^6^ CFU. This yielded the synTCS-MutLib strain, which comprises the total population of mutant DcuS/EnvZ chimeras. Flow cytometry analysis of synTCS-MutLib revealed a bimodal fluorescence distribution when grown in the absence of ligand, 20 mM fumarate, and 20 mM aspartate (Fig. 2A), indicating that mutagenesis had successfully altered the signaling outputs of mutant DcuS/EnvZ chimeras. We hypothesize that mutagenized DcuS/EnvZ chimeras have become constitutive or otherwise signaling-competent with potentially modified outputs and thus categorize these as loss-of-function (LOF) and gain-of-function (GOF), respectively.

**Figure 2.**
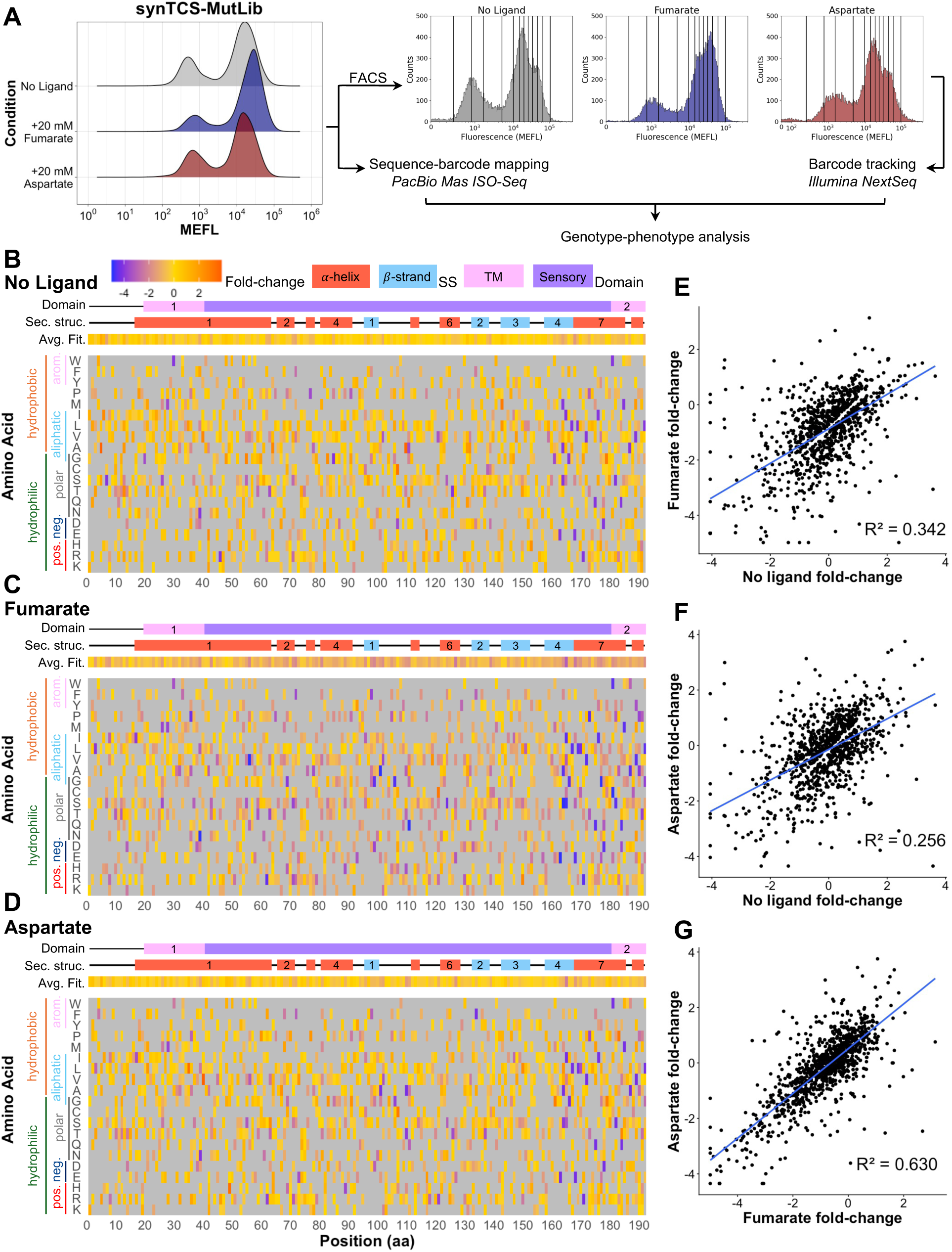
Mutational landscape of synTCS-MutLib. (A) Flow cytometry analysis of synTCS-MutLib grown without ligand, 20 mM fumarate, and 20 mM aspartate; Flow-seq enabled barcode tracking in synTCS-MutLib. Fold-change scores for 1,173 amino acid mutations in the DcuS sensory and TM domains for (B) no ligand, (C) 20 mM fumarate, and (D) 20 mM aspartate conditions. Scatter plots and correlations of (E) fumarate and no ligand, (F) aspartate and no ligand, and (G) fumarate and aspartate fold-change scores.

To understand sequence-function relationships within mutant chimeras in synTCS-MutLib, we needed to map the mutant sequences to their individual barcodes. The full DcuS/EnvZ coding sequence and downstream barcode region in synTCS-MutLib were PCR amplified and sequenced with PacBio Kinnex, yielding 818,700 amino acid sequences mapped to at least one barcode. To associate these mutant sequences with phenotypes, a Flow-seq (fluorescence-activated cell sorting and barcode tracking) approach was applied to different expression conditions. synTCS-MutLib was grown in three ligand conditions—no ligand, 20 mM fumarate, and 20 mM aspartate—and sorted into twelve bins of increasing fluorescence intensity (Fig. 2A). Sorted cells were cultured and barcodes were extracted via two-step PCR for Illumina NextSeq short-read sequencing. We filtered the sequence dataset to only include barcodes with ≥ 10 reads which were also successfully mapped to mutant sequences, resulting in 78%, 52%, and 70% of total sequencing reads being retained from the no ligand, fumarate, and aspartate datasets, respectively, and yielding 64,671 unique mapped barcodes.

To estimate a median activation level for each mutant, barcodes were first assigned to a “median bin” by performing a cumulative sum on normalized barcode frequencies across all twelve sorting bins for each condition. Then, median bins were converted to an inferred MEFL value by linear interpolation between the log_10_ of the bin boundaries in MEFL, which were measured prior to sorting (Table S2). A detailed flow chart of this pipeline is provided in Fig. S2. The unmutated DcuS/EnvZ sequence (wildtype) was observed for 3,957 barcodes, while the median barcode count for multi-barcoded mutant sequences was 32. The wildtype exhibited inferred fluorescence values of 13914 MEFL without ligand, 24888 MEFL with 20 mM fumarate, and 15970 MEFL with 20 mM aspartate. The wildtype-derived values were used to calculate fold-change scores (Eq. 2, Methods) for each mutant-barcode pair. Additionally, DNR and ligand specificity scores (Eq. 3, Methods) were calculated, resulting in six total phenotype scores for 64,671 full-length sequences.

### Mutational landscape of the DcuS sensory and transmembrane domains

To analyze the mutational landscape of the DcuS/EnvZ chimera, we used the fold-change score to quantify the effect of mutations on signaling outputs relative to wildtype. Heatmap visualization of fold-change scores for 1,173 amino acid mutations revealed several trends (Fig. 2B-D). On average, fumarate fold-change scores were more negative compared to those observed in the no ligand and aspartate conditions, indicating that, on average, mutations reduced fumarate-dependent signaling changes, which is expected when stochastically introducing mutations to a functional receptor.

Mutations associated with strongly negative fold-change scores across all three conditions clustered within residues 160-193, which correspond to a C-terminal 𝛽-strand and 𝛼-helix in the sensory domain and TM2 (Fig. 2B-D). This region is critical for transducing the signal from ligand binding from the periplasm to the cytosolic domains.^8,14^ In DcuS, ligand binding induces a compaction then uplift by the periplasmic sensory domain, which in turn pulls TM2 upwards into the periplasm and stimulates autokinase and phosphotransferase activity in the CA and DHp domains.^57^ We hypothesize that mutations in this region disrupted these conformational changes, resulting in loss of ligand-induced changes in signaling output. Within this region, we also observed decreased fumarate-activation for W181R (n = 20 barcodes; Supplementary Data 1), consistent with previously reported characterization of a DcuS^W181R^ mutant.^57^ Interestingly, the TM1 domain does not have a cluster of negative fold-change scores, suggesting it is more amenable to sequence diversification while maintaining function. TM1 does not significantly reposition during signal transduction; rather, it acts as a stator for TM2.^58^ Collectively, these data indicate that TM’s necessity to dynamically reposition limits its mutational tolerance.

To investigate how fold-change scores correlate within this dataset, we first filtered out barcode-sequence pairs with an aspartate and/or fumarate DNR between 0.1 and -0.1, which would encompass mutants with LOF phenotypes. These were removed to reduce bias from constitutive activity. Overall, fold-change scores were weakly correlated between the no ligand and ligand conditions (R^2^ = 0.342 for fumarate; 0.256 for aspartate), indicating that that many mutants have differing signaling outputs in the presence and absence of a ligand (Fig. 2E-F). A modest correlation (R^2^ = 0.630) was observed between the fumarate and aspartate conditions (Fig. 2G). Considering that mutations are distributed beyond the sensory domain, this partial covariation could arise from reweighting of ligand-dependent signaling outputs rather than altered ligand perception.

### Gain-of-function analysis reveals positions enriched for aspartate-dependent signaling outcomes

Having mapped global mutational effects, we next asked which positions enhanced aspartate-depending signaling. We performed a GOF analysis to determine which positions significantly altered aspartate responsiveness and, separately, specificity. We identified 634 single amino acid mutations with an aspartate DNR magnitude greater than the wildtype (0.19) and 291 single amino acid mutations with a ligand specificity score greater than 0. After filtering, the total number of observed barcodes at each position were summed. Barcode counts were used as a proxy for statistical power as each barcode represents an independent measurement. A total of eleven positions were enriched for both aspartate responsiveness and specificity, occurring more frequently based on a threshold of the mean plus two standard deviations of barcode counts across all positions (μ + 2σ; Fig. 3A-B).

**Figure 3.**
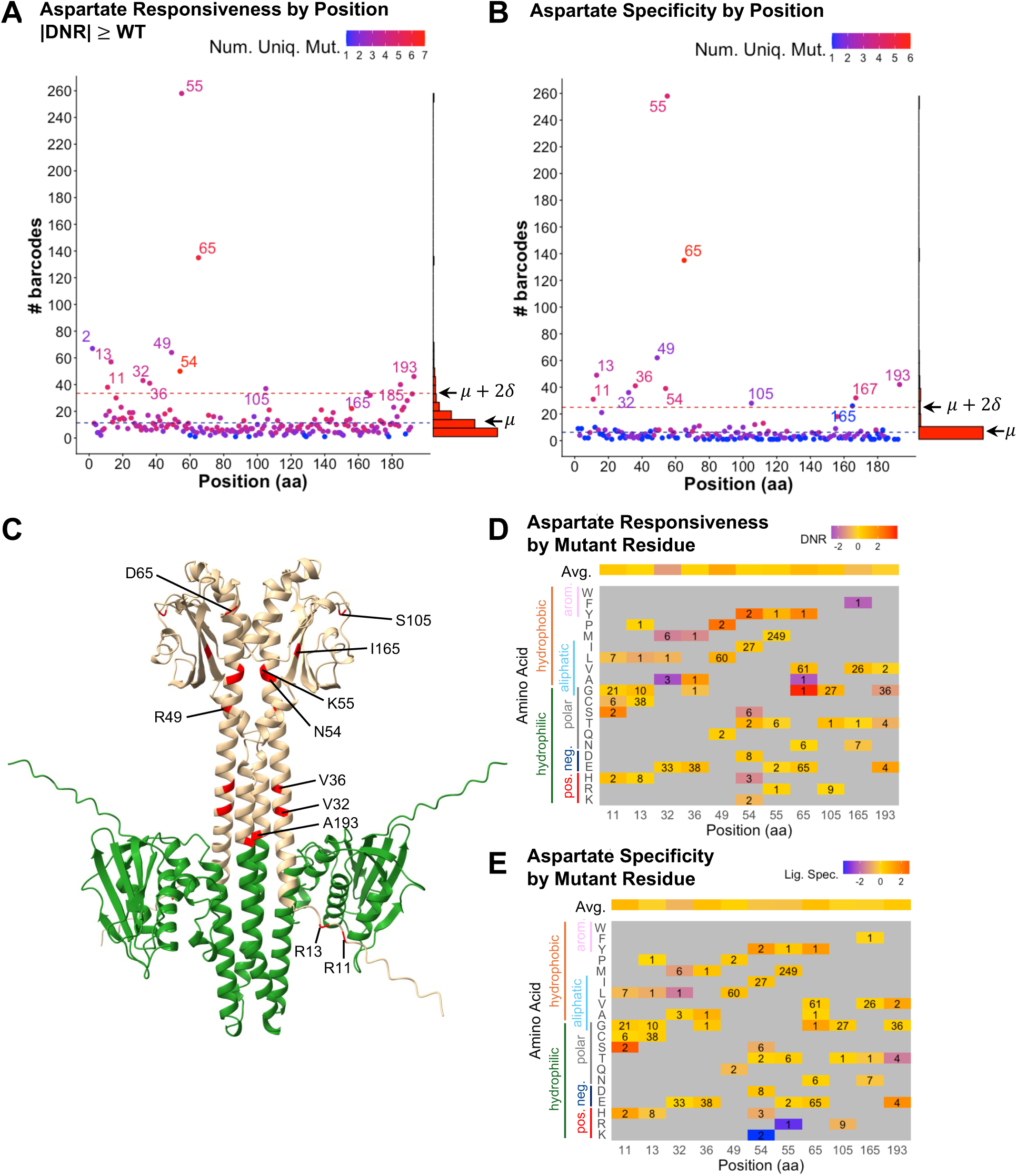
GOF analysis for aspartate responsiveness and specificity among single amino acid mutations in synTCS-MutLib. Eleven statistically significant positions were significantly enriched for both aspartate responsiveness (A) and specificity (B). Heatmap showing (C) aspartate DNR and (D) ligand specificity at each identified position. (E) Alphafold2 Multimer predicted structure of the DcuS (beige) EnvZ (green) chimera with positions of interest in red. (F) Correlation between fumarate and aspartate DNR values observed for the eleven significant positions, colored by ligand specificity score.

Interestingly, GOF positions clustered in two regions of the DcuS domain: the sensory domain and membrane-integrated regions of both TM domains (Fig. 3C). Mutations to TM domain residues V32, V36, and A193 tended to produce a neutral ligand specificity score, meaning that fumarate and aspartate signaling was similarly impacted (Fig. 3D-E). In contrast, mutations to sensory domain residues, R49, N54, K55, D65, S105, and I165, produced more diverse phenotypes (Fig. 3D-E). Within these positions, bulky aromatic amino acids, tryptophan and phenylalanine, frequently increased aspartate DNR, although this interpretation is limited due to the low barcode counts, and thus low confidence, observed for these mutations. In general, mutant amino acids with different chemical properties relative to the native residue tended to produce more substantial changes to phenotypes, indicating that disruption of local side chain interactions in the sensory domain tuned aspartate responsiveness and specificity.

Unexpectedly, two of the first fifteen residues (R11 and R13) were also found to significantly influence aspartate-dependent signaling outcomes. Currently, there are no reports of a functional role for this unstructured N-terminal region in DcuS. Due to their distal location from the ligand binding interface in the sensory domain^59,60^ and proximity to where TM1 is embedded in the plasma membrane,^58^ it is plausible that these positions instead are involved in tuning signaling outputs. Both positions identified were natively arginine residues, suggesting a potential importance of arginine’s chemical properties; however, further data are needed to elucidate the mechanism by which this region contributes to SHK signaling outputs.

Although some mutant positions show increased aspartate responsiveness and selectivity in comparison to wildtype, this does not directly translate to strong activation, as seen with wildtype fumarate responses (Fig. 1D, F). This is because wildtype aspartate DNR is minimal, so barcodes associated with modest activation were counted as significant for its mutant position. Indeed, few of the identified point mutations produced strongly increased aspartate responsiveness or selectivity. To develop DcuS-like SHKs with higher aspartate responsiveness, it may be necessary to perform targeted saturated mutagenesis on the identified positions and further explore the sequence space.

### Dial-out PCR enables individual mutant DcuS/EnvZ chimera characterization

To investigate mutant DcuS/EnvZ chimeras beyond our population-level Flow-seq analysis, dial-out PCR was used to isolate a subset of multi-barcoded mutants from synTCS-MutLib. Kruskal-Wallis and post-hoc Dunn tests were used to determine significant differences (p-value ≤ 0.05) between fold-change scores across all conditions from ≥ 3 barcodes for a given mutant. Mutant genes of interest were isolated from the library using barcode-specific primers, cloned into synTCS in place of the unmutated DcuS/EnvZ chimera, and individually characterized via flow cytometry.

Strikingly, mutant S4A produced a strongly activated constitutive phenotype, despite its distal location to the sensory domain (Fig. 4A). This result corroborates the conclusions from our GOF analysis that positions in the first fifteen amino acids of DcuS/EnvZ play a functional role in influencing signaling outputs. Although position 4 itself was not identified as significant for driving GOF for aspartate-dependent signaling outcomes, that residues in this region of DcuS was implicated in adjacent significance analyses implies biologically relevant function for SHK signaling.

**Figure 4.**
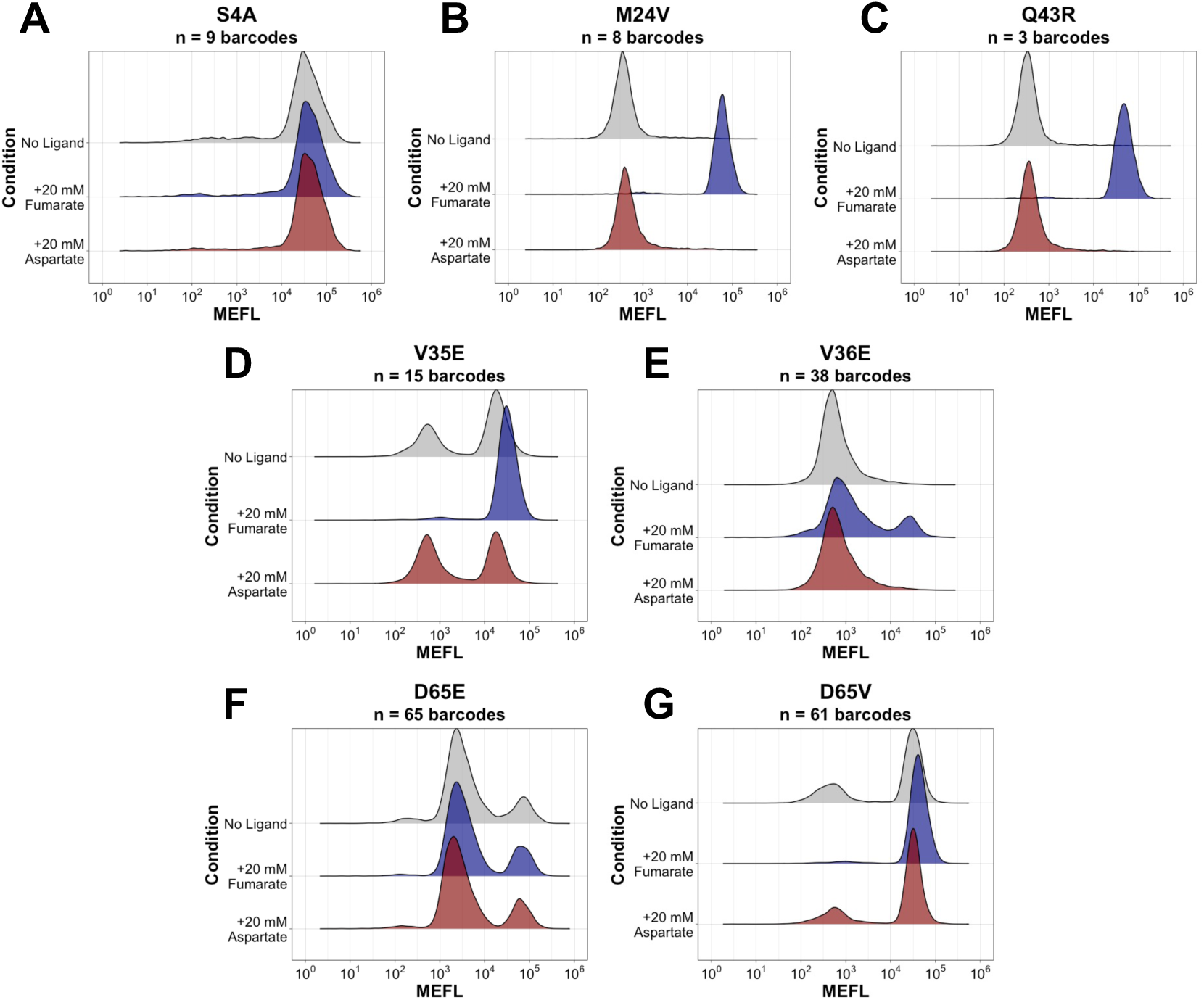
Flow cytometry analysis of seven dial-out mutants: (A) synTCS-S4A, (B) synTCS-M24V, (C) synTCS-Q43R, (D) synTCS-V35E, (E) synTCS-V36E, (F) synTCS-D65E, and (G) synTCS-D65V.

Two mutants which retained ligand responsiveness, mutants M24V and Q43R, did not significantly alter the chimera’s fumarate or aspartate responsiveness relative to the wildtype (Fig. 4B-C), similar to our findings that TM1 mutations produced fewer strongly negative or positive fold-change scores compared to TM2 (Fig. 2E-G). To further understand the role of mutations in TM1, mutants V35E and V36E mutants were characterized and found to produce distinctive LOF phenotypes (Fig. 4D-E). While mutant V35E retained fumarate responsiveness, it exhibited bimodal distribution in non-activating conditions, suggesting that the mutation disfavors the ‘off’ state of the sensor. Contrastingly, mutant V36E exhibited an opposite activation profile: a low fluorescence population was observed in all conditions, with a small subpopulation of highly fluorescent cells when exposed to fumarate. Despite the shared chemical properties of native and mutant residues, their position in the TM1 helix influenced the preferred signaling state of the mutant chimera, indicating that a residue’s location within the chimera determines its role in mediating signal processing.

We further investigated biochemically distinct mutations at the same position: mutant D65E produced an intermediate constitutively active phenotype, while mutant D65V produced a stronger constitutively active phenotype (Fig. 4F-G). Although there are similarities between the resultant phenotypes of these mutants that of mutants V35E and V36E, the mechanisms by which they arise may be differentiated by the domain in which they are found. Due to their proximity to the ligand binding interface in the sensory domain,^59,60^ mutants D65E and D65V may instead influence signaling by interfering with ligand perception. One plausible explanation for this is that side chain interactions within the sensory domain may be impacted by these mutations, although further structural analyses are required.

One limitation of this approach lies in the accuracy of inferred MEFL values when compared to individually characterized phenotypes. Across all multi-barcoded mutants observed in the library, the standard deviation of inferred MEFL for all conditions sharply decreases as barcode count increases (Fig. S3). This variability is due to the inherent noise associated with each barcode measurement, which may arise from sorting stochasticity, growth differences, and PCR bias during barcode extraction. One caveat is that barcode counts less than twenty were also observed to have some of the lowest standard deviations between reads, tending to occur in mutants with 2 barcodes; however, this may reflect the smaller sample size rather than true measurement consistency. Furthermore, mutants with bimodal responsiveness also correlated less with their predicted phenotypes, highlighting the importance of utilizing a higher barcoding threshold to comprehensively and accurately sample a given mutant’s fluorescent phenotype in pooled high-throughput assays.

## Conclusions

Here, we have developed and applied a synthetic TCS platform for characterizing the mutational landscape of the C4-dicarboxylate receptor, DcuS, in a chimeric context. By linking sequence-function relationships across 1,173 DcuS/EnvZ mutants, we identified several biologically relevant trends in fold-change of mutations to the DcuS sensory and TM domains relative to the wildtype. Importantly, we found that mutations in the TM2 domain frequently drove strongly negative fold-change scores while TM1 mutations did not produce similar trends, which may be related to the roles of each domain in signal transduction. Additionally, our GOF analysis revealed two residues in the first fifteen N-terminal positions of DcuS to be functionally relevant for attenuating aspartate-dependent signaling and individual characterization of a dialed-out S4A mutant confirmed the role of this region in influencing signaling output of the DcuS/EnvZ chimera. Although scaling this approach in multiplexed assays remains challenging to apply to phenotypically diverse SHKs, constructing a library wherein sequences are associated with a higher number of barcode identifiers to improve sample size in pooled assays would greatly improve measurement accuracy. In future, libraries of rewired chimeric SHKs may be screened for responsiveness towards many different metabolites and expedite the discovery and creation of novel biosensors.

## Methods

### Cloning

A three-fragment Golden Gate assembly was used to clone the DcuS/EnvZ chimera into a plasmid vector. The first fragment (FrgA), comprising the TM1, sensory, and TM2 domains (amino acids 1-193), was amplified from the genome of E. coli K12 strain MG1655 using primers found in Table S1. The second fragment (FrgB) consisted of the CA and DHp domains (amino acids 232-451) of EnvZ and was amplified from the genome of E. coli K12 strain MG1655 using primers found in SI Table 1. Additionally, a 24-base pair (bp) quasi-randomer barcode region (5’-NNBBDDBBVVHHDDBBVVHNDDNN-3’) was inserted into FrgB via PCR. The final fragment (FrgC) was derived from the pHKGG1 vector and contained a p15A Ori, carbenicillin selection marker, lac repressor, and P_trc_ inducible promoter; FrgA and FrgB were cloned downstream of the P_trc_ promoter.

Four Golden Gate reactions were run using the following temperature program: 1) incubate at 37°C for 20 hours; 2) heat inactivation of both enzymes at 80°C for 20 mins; 3) final hold at 12°C. The molar ratios used were 0.18 pmol for FrgC, 0.36 pmol for FrgB, and 0.36 pmol for FrgA (DcuS sensory domain) for a total of 470 ng of DNA in the reaction. The reagent volumes were 2.5 𝜇L of 10x T4 DNA Ligase Buffer (NEB), 2.5 𝜇L of 10 mM ATP (NEB), 0.25 𝜇L T4 DNA Ligase (NEB), 0.75 𝜇L BsaI-HFv2 (NEB), and water to complete the remaining volume to a total of 25 𝜇L. Assemblies were then pooled and cleaned using the Monarch PCR and DNA Cleanup Kit (NEB) and then drop dialyzed on a 0.05 𝜇m membrane filter (Sigma Millipore) for a minimum of 30 minutes. Purified pDcuSEnvZ was used as the input for electroporations (Bio-Rad MicroPulser), which were then combined and plated.

### Integration of synthetic two-component signaling machinery

The pSR40.29 plasmid, harboring chimeric response regulator OmpR/CcrA and superfolder GFP (sfGFP),^50^ was transformed via electroporation into E. coli strain BW29655, which lacks the endogenous *envZ* and *ompR* genes in its genome.^51^ The resultant strain, pSR, was made electrocompetent by repeated steps of centrifugation at 4,000 g for 15 minutes at 4°C and resuspension of bacterial cells in 50mL, 25mL, 10mL, and 1mL of ice-cold 10% glycerol. The purified pDcuSEnvZ was then transformed via electroporation into pSR, resulting in the strain synTCS.

### Mutagenesis of the DcuS sensory and transmembrane domains

Error prone PCR with a GeneMorph II Random Mutagenesis Kit (Agilent) was used to introduce random mutations to amino acid residues 2-193 of *dcuS*, which comprises the TM1, sensory, and TM2 domains. We followed the recommended protocol (Agilent) to achieve a mutation rate of 0.4-5 per kilobase pair (kbp). Cloning occurred the same as the unmutated DcuS/EnvZ chimera. The resultant purified mutagenic library plasmids were first transformed via electroporation (Bio-Rad MicroPulser) into electrocompetent E. coli 10𝛽 (NEB) for initial cloning then into electrocompetent pSR. Serial dilutions were used to determine the library diversity. The mutagenic library as a whole population is henceforth referred to as strain synTCS-MutLib.

### Homemade supplemented M9 minimal medium

Supplemented M9 minimal medium (M9MM) was made and used throughout this study for microwell plate, flow cytometry, and fluorescence-activated cell sorting experiments. 1×M9 salts, 2 mM MgSO_4_, 0.1 mM CaCl_2_, 0.4% (wt/vol) galactose, 0.2% (wt/vol) casamino acids (Gibco) were combined. When appropriate, 1X chloramphenicol (CAM) and carbenicillin (Carb) were added for plasmid maintenance; 100 ng/mL anhydrotetracycline (aTc; Takara Biosciences) and 1 mM isopropyl β-D-1-thiogalactopyranoside (IPTG; Thermo Scientific) for *ompR/ccrA* and *dcuS/envZ* expression induction, respectively; and 3-20 mM disodium fumarate or 3-20 mM aspartic acid for ligand responsiveness. All reagents were supplied by Sigma Aldrich unless otherwise specified.

### Time-course microwell plate fluorescence assays

96-well plates (Invitrogen M33089) were inoculated with supplemented M9MM, cell culture to a final OD_600_ of 0.1, and appropriate antibiotics and inducers. A 12-step serial dilution of 0.017 𝜇M fluorescein was used as a green fluorescence control.^61^ Plates were grown for twelve hours shaking linearly at 37°C. The BioTek Synergy H1 (Agilent) plate reader and the Gen5 3.14 software (Agilent) were used to measure the OD_600_ and fluorescence (excitation 𝜆: 485 nm; emission 𝜆 : 515 nm) every 6 minutes. The resultant data was analyzed using tools from wellplate_analysis.^62^ Fluorescence was converted to MEFL using a linear regression derived from the fluorescein calibration curve, then divided by OD_600_ to normalize.

### Culture preparation for flow cytometry and fluorescence-activated cell sorting

synTCS was inoculated in 5 mL of homemade M9 minimal media supplemented with 100 ng/mL aTc, 1 mM IPTG, 1X CAM, 1X Carb, and 3, 10, or 20 mM fumarate or aspartate. synTCS-MutLib was inoculated at 0.1 OD_600_ in 40mL homemade M9 minimal media supplemented with 100 ng/mL aTc, 100 mM IPTG, 1X CAM, 1X Carb, and 20 mM of either fumarate or aspartate. As controls, BW29655 without any plasmid (WT) was growth in 5mL M9MM. After 6 hours of growth shaking at 37°C, 5mL of each culture (25mL for synTCS-MutLib) was centrifuged at 4,000 g for 5 minutes at 4°C. After decanting the media and resuspending the pellet in 1X phosphate buffered saline (PBS), the cell solution was strained through a 20 μM filter (pluriSelect) and diluted to 7×10^7^ cells/mL in 1X PBS. Additionally, Rainbow Calibration beads (Spherotech, RCP-30-5A) were added to 2 mL 1X PBS following manufacturer’s guidelines and run prior to sample acquisition.

### Flow cytometry analysis

The BD Accuri cell sorter was used to analyze all samples. Sample acquisition occurred at an event rate of < 10,000 events/second. All samples were gated for singlets in FlowJo 11 (Fig S4), fluorescence values converted from flow cytometry fluorescence units to molecules of equivalent fluorescein (MEF) in FlowCal^63^ using Rainbow Calibration beads (Spherotech, RCP-30-5A), and visualized with built-in functions from FlowCal in Python or ggplot2 in R (4.4.1).

### Fluorescence-activated cell sorting of synTCS-MutLib

Preparation of synTCS-MutLib cultures was consistent with all flow cytometry experiments, except a culture volume of 150 mL was used for sorting. The BD Symphony S6 was used with a 70 𝜇m nozzle and BD FACSDiva (BD Biosciences) software for fluorescence-activated cell sorting (FACS). Singlets were selected for by gating in the FSC detection range, and sample binning occurred using green fluorescence. synTCS-MutLib grown with 20 mM fumarate, 20 mM aspartate, and without ligand was sorted into twelve bins; bin boundaries were determined based on prior flow cytometry analysis of 100,000 cells from the mutagenic library (Fig. 2A). Flow cytometer fluorescence units and converted MEFL values for bin boundaries are reported in Table S2. Samples were processed and analyzed in the same manner as flow cytometry samples. Sample acquisition was carried out at < 18,000 events/second. Approximately 800,000 events were collected and sorted for each bin. All cells were sorted into epi tubes filled with 300 μL Lauria-Bertani (LB; BD Biosciences) broth, placed on ice, then plated on 100mm × 100mm agar plates (Corning) with appropriate antibiotics. After 16 hours of growth at 37°C, colonies were scraped, combined with 50% glycerol, and stored at -80°C.

### MAS Iso-seq and analysis of barcode-sequence pairs in mutagenic library

synTCS-MutLib was PCR amplified with HK_PB_03_FWD/REV primer pair (Table S1) using Q5 DNA polymerase and 10 ng of template (miniprepped plasmid) in a 50 μL reaction for 12 cycles. The resultant synTCS-MutLib PCR product was submitted to the UOregon Genomics and Cell Characterization Core Facility (GC3F) core for library preparation and sequencing by MAS ISO-seq (Kinnex). Raw PacBio variant-barcode mapping reads were submitted to the NCBI Sequence Read Archive under BioProject accession PRJNA1468850. For the MAS ISO-seq data, skera (0.1.0) was used to split the MAS arrays within 1,268,200 CCS reads, producing 15,907,686 split segments. Lima^64^ was then used to demultiplex the split segments, resulting in 2,740,434 to 4,826,815 reads per library. For MAS ISO-seq data, we first identified the constant regions flanking the barcode (GTCGCTGCCGAACAGC-24N-AGGAGAAGAGCGCACG), allowing up to 3 mismatches. Then each read was scanned for the presence of the NdeI (CATATG) site at the start codon and the conserved region in the vector immediately flanking the cloning site (GACGACCGCACGCTGCTG, which corresponds to residues 232-237 (DDRTLL) of EnvZ). The script outputs this variable region and each associated barcode, as well as the barcodes counts. Barcode counts were inputted into Starcode (1.4)^65^ for collapse with a distance of 1 using the sphere algorithm. A consensus call was made for each barcode using abPOA.^66^ Genes were aligned to the DcuS wildtype sequence using minimap2,^67^ with k-mers set to 10. Scripts for the full pipeline are available in our GitHub repository.^68^ We found that only 3% of the sequenced mutagenic library encode full length DcuS TM1, sensory, and TM2 domains, indicated an unexpectedly high rate of mutations that resulted in truncated products; we suggest that future studies use a lower mutagenesis rate or nicking mutagenesis approaches instead.

### Illumina NextSeq and analysis of barcodes obtained from fluorescence-activated cell sorting bins

The sorted synTCS-MutLib samples were miniprepped (ThermoScientific GeneJet Plasmid Miniprep Kit) and PCR amplified with a mix of five primer pairs, Illumina_adapter_camp1-5_fwd/rev (Table S1), using Kapa HiFi DNA polymerase (NEB) and 4 ng of template in a 25 μL reactions for 16 cycles. The resultant 98-108 bp products were size selected, PCR amplified using one of 36 primer pairs (Table S1) with one primer pair associated with each bin in all three conditions, using Equinox DNA polymerase (Watchmaker Genomics) and extremely thermostable single stranded DNA binding protein (ETSSB; NEB) in 50 μL reactions for 10 cycles. The resultant 218-228 product was size selected. This process was repeated on the original pMutLib prior to transformation into pSR (presort library 1), and pMutLib miniprepped from synTCS-MutLib prior to sorting (presort library 2). All 38 of these library samples were mixed into a single 75 μL sample containing 20 nM of DNA per sample and submitted to the UOregon GC3F core for library preparation and sequencing with Illumina NextSeq P4000 paired ends. Raw Illumina barcode reads for all conditions and FACS bins were submitted to the NCBI Sequence Read Archive under BioProject accession PRJNA1468850.

Demultiplexed FASTQ files were processed with an in–house workflow.^68^ BBMerge^69^ from BBTools was used to merge overlapping paired reads into a single read. Next, hts_primers^70^ from HTStream was used to flip the sequences into the correct orientation and trim the primer sequence from the ends of the sequences. The quality data was removed to turn the FASTQ files into FASTA format, and the first 24 bases (the length of the barcode on the SHK plasmids) were cut out, with the number of times each barcode appeared counted. Starcode,^65^ a DNA sequence clustering software, was then used to cluster and collapse all barcodes and their counts at a Levenshtein distance of one for the final output of this initial pipeline. Processing was performed independently for every library–bin combination, yielding barcode–count matrices that fed subsequent activity inference analyses.

### Association of mutant sequence, barcode, and inferred fluorescence phenotype

The total set of barcodes across all bins in the three conditions—no ligand, fumarate, and aspartate—were filtered based the following criteria: if a barcode had been mapped to a mutant sequence in the PacBio Mas ISO-Seq analysis and had more than 10 reads associated with it. Using matrices of barcode counts in each bin for all three conditions, the protocol from Biswas et al. (2021)^71^ was adapted to infer median bin scores for each barcode in all three conditions. First, the relative abundance of all barcodes in each bin was computed by dividing barcode reads by the total reads in that bin. Next, the fold-change abundances of barcodes in each bin were computed by dividing the relative abundances by the abundance of that barcode in presort library 2. Third, the frequency each barcode appears in each bin was calculated by dividing the fold-change abundances of that barcode in each bin by the sum of the fold change abundances of the barcode across all bins. A cumulative sum of frequencies was performed, and a median bin was assigned to each barcode at a cumulative sum of 0.5. These median bin scores were converted to molecules of equivalent fluorescein (MEFL) by linear interpolation between the log_10_ of MEFL values of the bin boundaries flanking each median bin score.

Phenotype scores for each mutant-barcode pair were calculated using equations (1), (2), and (3) using wildtype and mutant-specific inferred MEFL values for each condition. This yielded six phenotype scores for each mutant: no ligand fold-change, fumarate fold-change, aspartate fold-change, fumarate dynamic range, and aspartate dynamic range, and ligand specificity.

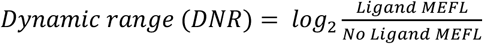

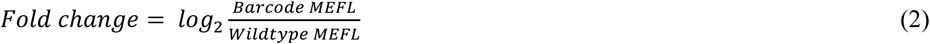

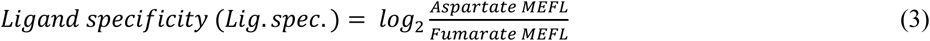

All scores were assigned to amino acid mutations by identifying differences in wildtype and mutant sequences. For mutations with more than one set of phenotype scores, the average, standard deviation, and 95% confidence interval of all scores were calculated for each. This pipeline is summarized in Fig. 6; it was performed in R (4.4.1) and is accessible in our GitHub repository: https://github.com/PlesaLab/DcuSEnvZ.

### Structural annotations for heatmaps

Secondary structure annotations were generated using the Dictionary of Secondary Structure in Proteins (DSSP) server.^72–73^ Domain annotations for the DcuS TM and sensory domains were adapted from previously annotated DcuS structures (PDB: P0AEC8).

### Gain-of-function analysis

To identify positions which significantly influence aspartate responsiveness and specificity, the resultant Flow-seq dataset (see: Association of mutant sequence, barcode, and inferred fluorescence phenotype) was filtered then collapsed by position. For aspartate responsiveness, only barcode-sequence pairs with an absolute value of aspartate DNR greater than the wildtype aspartate DNR in the dataset (0.19) were kept; for aspartate specificity, only barcode-sequence pairs with a ligand specificity score greater than 0 were kept. Then, the filtered datasets were collapsed by position, and the total number of unique mutant amino acid residues and barcodes were summed. The average and standard deviation of barcode counts were calculated, and a position was considered significant if its barcode count was above two standard deviations plus the mean for observed barcode counts in each filtered dataset. Positions that were enriched in both datasets were considered to be significant position determinants of aspartate responsiveness and selectivity.

### Statistical analysis of multi-barcoded mutant sequences

To identify DcuS/EnvZ mutants with altered signaling behavior, we filtered for mutant sequences that are associated with at least three barcodes. Kruskal-Wallis and post-hoc Dunn tests were performed to identify mutants with median MEFL fluorescence and median fold-change scores that differ significantly across at least two of the three conditions. This analysis was performed in R (4.4.1) and is accessible in our GitHub repository: https://github.com/PlesaLab/DcuSEnvZ. The resultant 33 mutants that were identified as significant were selected for wet lab validation.

### Machine learning predictions of mutational fitness effects in DcuS/EnvZ chimera

Computational predictions of mutational effects were generated using VenusREM,^74^ a state-of-the-art zero-shot protein fitness model that integrates sequence, structure, and evolutionary context. The wildtype amino-acid sequence of the DcuS N-terminal portion of the chimera was provided as input (RHSLPYRMLRKRPMKLSTTVILMVSAVLFSVLLVVHLIY FSQISDMTRDGLANKALAVARTLADSPEIRQGLQKKPQESGIQAIAEAVRKRNDLLFIVVTDMQSLRYSHPEAQRIGQPFKGDDILKALNGEENVAINRGFLAQALRVFTPIYDENHKQIGVVAIGLELSRVTQQINDSRWSLQMAAGVKQLA). A structural homodimer model of the DcuS region was obtained using AlphaFold3.^75^ Only chain A of the predicted structure was used for downstream analysis. Structural tokens were derived from the AlphaFold3 model using the ProSST discretization procedure implemented in the VenusREM pipeline. A multiple sequence alignment (MSA) was generated automatically using the EVcouplings^76^/JackHMMER^77^ workflow. For each possible single amino acid substitution, VenusREM computed a log-likelihood ratio score reflecting the model estimated change in fitness relative to wild type. These scores were obtained using the default VenusREM settings (ProSST-2048 backbone with MSA-based retrieval). Model predictions were then compared directly to our experimental fumarate DNR scores, however, no correlation was found (Fig. S5).

### Validation of significantly performing multi-barcoded mutants

To validate mutants individually, the top two most abundant barcodes in the presort library 2 data were used to design primers which are specific to each mutant for dial-out PCR. Primers were designed using a custom Python script, primer_design_MP.py, which can be accessed at our GitHub repository. Size-selected pDcuSEnvZ plasmids were miniprepped from unsorted synTCS-MutLib samples then PCR amplified using mutant-specific primers (Table S1) with Equinox DNA polymerase (Watchmaker Genomics), ETSSB (NEB), and 1 ng of template in 50 𝜇L reactions for 10 cycles of touchdown and 35 additional cycles at the target annealing temperature. The resultant 1.3 kbp products were digested with NdeI and KnpI and ligated with gel-extracted FrgC, then transformed into electrocompetent pSR. Three colonies per mutant were sequenced, and successful hits were analyzed using the above protocols for flow cytometry assays.

## Conflicts of Interest

C.P. holds equity in SynPlexity.

## Acknowledgements

This work was supported by a Career Award at the Scientific Interface from the Burroughs Wellcome Fund. L.L. was supported by the Knight Campus Undergraduate Scholars Program and the Undergraduate Research Opportunities Program Mini-Grant. We thank the UOregon GC3F core staff for help with sequencing. Plasmid pSR40.29 was a gift from Jeffrey Tabor (Addgene plasmid # 125088; http://n2t.net/addgene:125088 ; RRID:Addgene_125088). We also thank Alex J. Eddins, Anissa Benabbas, and Sayandeep Gupta for the many insightful conversations over the duration of this work.

## Abbreviations

TCS: Two-component signaling
SHK: Sensor histidine kinase
RR: Response regulator
TM: Transmembrane
DNR: Dynamic range
MEFL: Molecules of equivalent fluorescein
FACS: Fluorescence activated cell sorting

## Notes

https://github.com/PlesaLab/DcuSEnvZ

https://www.ncbi.nlm.nih.gov/bioproject/1468850

https://figshare.com/articles/dataset/DcuSEnvZ_Output_Data/32340699

