## Supplemental Information for "Characterizing Sequence-Function Relationships in Chimeric DcuS/EnvZ Histidine Kinases at Scale"

*Engineering Novel Signaling Phenotypes in Chimeric DcuS/EnvZ Histidine Kinases*

SUPPLEMENTARY FIGURES	page 2

**Figure S1.** Fluorescence over time of uninduced pSR and synTCS strains.

**Figure S2.** Overview of computational pipeline used to preprocess raw Illumina NextSeq data, infer MEFL fluorescence, and compute phenotype scores for all mapped sequence-barcode pairs.

**Figure S3.** Standard deviation of inferred MEFL values for mutants of varying barcode counts.

**Figure S4.** Example of singlets gating strategy for flow cytometry data.

**Figure S5.** Comparison of VenusREM-predicted and observed fitness scores.

SUPPLEMENTARY TABLES	page 5

**Table S1.** List of primers used in this study.

**Table S2.** Fluorescence-activated cell sorting bin boundaries in relative fluorescence units (RFU) and molecules of equivalent fluorescein (MEFL).

**Table S3.** Phenotype scores of all observed multi-barcoded mutants.

PLASMID MAPS AND SEQUENCES

SUPPLEMENTARY FIGURES

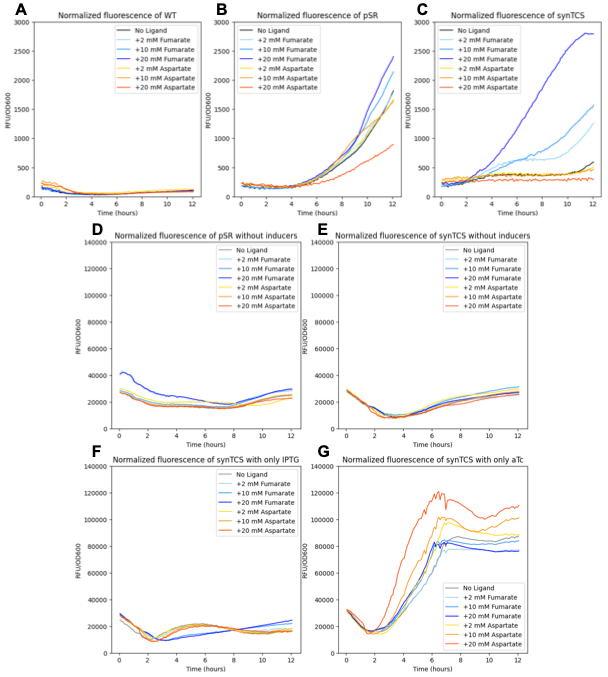

Figure S1. Normalized fluorescence over time of (A) WT, (B) pSR with aTc, (C) synTCS with aTc and IPTG, (D) pSR without inducers, (E) synTCS without inducers, (F) synTCS with only IPTG, and (G) synTCS with only aTc grown without ligand or 2, 10, or 20 mM fumarate or aspartate. The data shown in A, B, and C used different gain settings than the experiment in D, E, F, and G, and scales are not comparable.

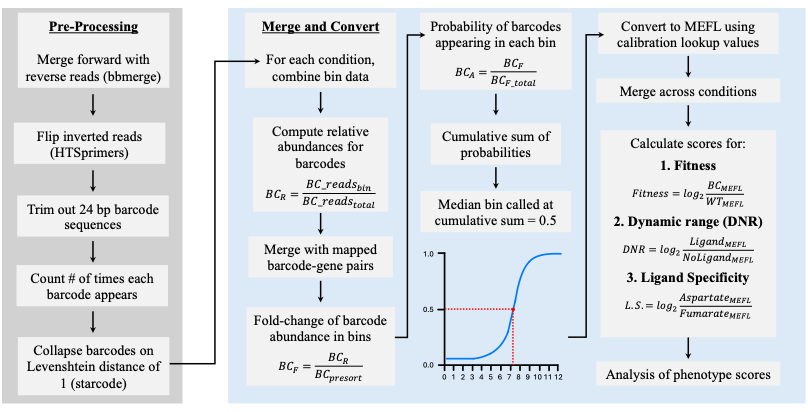
Figure S2. Overview of computational pipeline used to preprocess raw Illumina NextSeq data, infer MEFL fluorescence, and compute phenotype scores for all mapped sequence-barcode pairs.

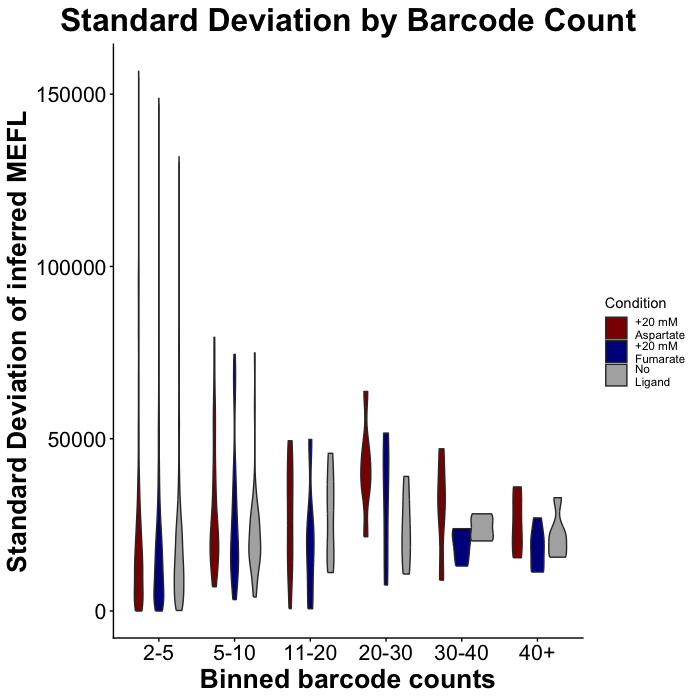

Figure S3. Standard deviation of inferred MEFL values for mutants of varying barcode counts.

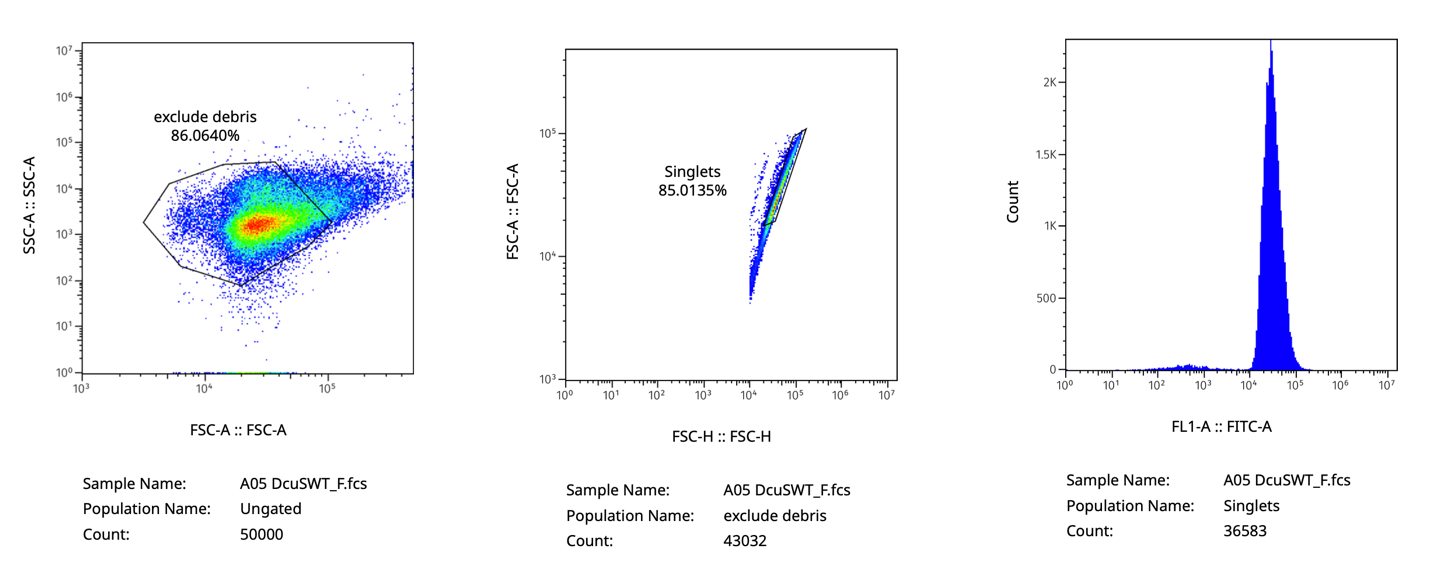

Figure S4. Example of singlets gating strategy for flow cytometry data.

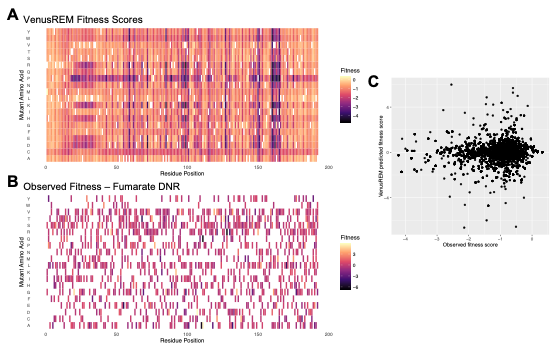

Figure S5. Comparison of VenusREM-predicted and observed fitness scores. Heatmaps of amino acid by position colored by (A) VenusREM scores and (B) observed fumarate DNR values. (C) Scatter plot of VenusREM-predicted and observed fitness scores.

SUPPLEMENTARY TABLES

Table S1. List of primers used in this study.

| **Primer name** | **Sequence (5'-3')** |
| --- | --- |
| DcuS_Ecoli_genome_fwd | tgcGGTCTCACATatgAGACATTCATTGCCCTACCGCA |
| DcuS_Ecoli_genome_rev | acgGGTCTCACGTCCGCCAGTTGCTTAACACCAGCCGCCATCTGCAGGCTCCAGCGACTGTCATTGATCTG |
| EnvZ_Ecoli_genome_fwd | gccgaaaccctggcgaaactgttta |
| EnvZ_Ecoli_genome_rev | aaccaggggcgttttcatctcgttg |
| HK_PB_03_FWD | CTACACGACGCTCTTCCGATCTACACATCTCGTGAGAGCACACAGCACTCTCCTAATTTAAATCGCACAACTCATATG |
| HK_PB_03_REV | AAGCAGTGGTATCAACGCAGAGCTCTCACGAGATGTGTCGTGACGTCGTGCGCTCTTCT |
| Illumina_adapter_camp1_fwd | GCTCTTCCGATCTNCCTAAGTGTCGCTGCCGAACAGC |
| Illumina_adapter_camp1_rev | GCTCTTCCGATCTNTGACGTCGTGCGCTCTTCTCCT |
| Illumina_adapter_camp2_fwd | GCTCTTCCGATCTNNCCTAAGTGTCGCTGCCGAACAGC |
| Illumina_adapter_camp2_rev | GCTCTTCCGATCTNNTGACGTCGTGCGCTCTTCTCCT |
| Illumina_adapter_camp3_fwd | GCTCTTCCGATCTNNNCCTAAGTGTCGCTGCCGAACAGC |
| Illumina_adapter_camp3_rev | GCTCTTCCGATCTNNNTGACGTCGTGCGCTCTTCTCCT |
| Illumina_adapter_camp4_fwd | GCTCTTCCGATCTNNNNCCTAAGTGTCGCTGCCGAACAGC |
| Illumina_adapter_camp4_rev | GCTCTTCCGATCTNNNNTGACGTCGTGCGCTCTTCTCCT |
| Illumina_adapter_camp5_fwd | GCTCTTCCGATCTNNNNNCCTAAGTGTCGCTGCCGAACAGC |
| Illumina_adapter_camp5_rev | GCTCTTCCGATCTNNNNNTGACGTCGTGCGCTCTTCTCCT |
| DcuSML_FACS_bin-NL1_D01_fwd | AATGATACGGCGACCACCGAGATCTACACCGACTCACTAACACTCTTTCCCTACACGACGCTCTTCCGATC*T |
| DcuSML_FACS_bin-NL1_D01_rev | CAAGCAGAAGACGGCATACGAGATTACACATAGGGTGACTGGAGTTCAGACGTGTGCTCTTCCGATC*T |
| DcuSML_FACS_bin-NL2_D02_fwd | AATGATACGGCGACCACCGAGATCTACACTTCCGTCCAAACACTCTTTCCCTACACGACGCTCTTCCGATC*T |
| DcuSML_FACS_bin-NL2_D02_rev | CAAGCAGAAGACGGCATACGAGATGTCAACGTTGGTGACTGGAGTTCAGACGTGTGCTCTTCCGATC*T |
| DcuSML_FACS_bin-NL3_D03_fwd | AATGATACGGCGACCACCGAGATCTACACATCTCTGGTAACACTCTTTCCCTACACGACGCTCTTCCGATC*T |
| DcuSML_FACS_bin-NL3_D03_rev | CAAGCAGAAGACGGCATACGAGATACCTATGTCAGTGACTGGAGTTCAGACGTGTGCTCTTCCGATC*T |
| DcuSML_FACS_bin-NL4_D04_fwd | AATGATACGGCGACCACCGAGATCTACACCCTAACTCATACACTCTTTCCCTACACGACGCTCTTCCGATC*T |
| DcuSML_FACS_bin-NL4_D04_rev | CAAGCAGAAGACGGCATACGAGATGTTGGTGTAAGTGACTGGAGTTCAGACGTGTGCTCTTCCGATC*T |
| DcuSML_FACS_bin-NL5_D05_fwd | AATGATACGGCGACCACCGAGATCTACACTTCGATGACGACACTCTTTCCCTACACGACGCTCTTCCGATC*T |
| DcuSML_FACS_bin-NL5_D05_rev | CAAGCAGAAGACGGCATACGAGATTAATTCCAGGGTGACTGGAGTTCAGACGTGTGCTCTTCCGATC*T |
| DcuSML_FACS_bin-NL6_D06_fwd | AATGATACGGCGACCACCGAGATCTACACAACTAGGTCAACACTCTTTCCCTACACGACGCTCTTCCGATC*T |
| DcuSML_FACS_bin-NL6_D06_rev | CAAGCAGAAGACGGCATACGAGATTTGATGTTGCGTGACTGGAGTTCAGACGTGTGCTCTTCCGATC*T |
| DcuSML_FACS_bin-NL7_D07_fwd | AATGATACGGCGACCACCGAGATCTACACAGTACCATGAACACTCTTTCCCTACACGACGCTCTTCCGATC*T |
| DcuSML_FACS_bin-NL7_D07_rev | CAAGCAGAAGACGGCATACGAGATACCTGGCTGTGTGACTGGAGTTCAGACGTGTGCTCTTCCGATC*T |
| DcuSML_FACS_bin-NL8_D08_fwd | AATGATACGGCGACCACCGAGATCTACACCTGTCCTGTCACACTCTTTCCCTACACGACGCTCTTCCGATC*T |
| DcuSML_FACS_bin-NL8_D08_rev | CAAGCAGAAGACGGCATACGAGATAGATGGACTAGTGACTGGAGTTCAGACGTGTGCTCTTCCGATC*T |
| DcuSML_FACS_bin-NL9_D09_fwd | AATGATACGGCGACCACCGAGATCTACACAGTCGTCACGACACTCTTTCCCTACACGACGCTCTTCCGATC*T |
| DcuSML_FACS_bin-NL9_D09_rev | CAAGCAGAAGACGGCATACGAGATTCTGAGAGGCGTGACTGGAGTTCAGACGTGTGCTCTTCCGATC*T |
| DcuSML_FACS_bin-NL10_D10_fwd | AATGATACGGCGACCACCGAGATCTACACGCCGGTAAGTACACTCTTTCCCTACACGACGCTCTTCCGATC*T |
| DcuSML_FACS_bin-NL10_D10_rev | CAAGCAGAAGACGGCATACGAGATCTTGAACCTAGTGACTGGAGTTCAGACGTGTGCTCTTCCGATC*T |
| DcuSML_FACS_bin-NL11_D11_fwd | AATGATACGGCGACCACCGAGATCTACACGAAGGAGAGCACACTCTTTCCCTACACGACGCTCTTCCGATC*T |
| DcuSML_FACS_bin-NL11_D11_rev | CAAGCAGAAGACGGCATACGAGATCGAGAGTCTTGTGACTGGAGTTCAGACGTGTGCTCTTCCGATC*T |
| DcuSML_FACS_bin-NL12_D12_fwd | AATGATACGGCGACCACCGAGATCTACACCGTGGTCACTACACTCTTTCCCTACACGACGCTCTTCCGATC*T |
| DcuSML_FACS_bin-NL12_D12_rev | CAAGCAGAAGACGGCATACGAGATGTCGATCTTCGTGACTGGAGTTCAGACGTGTGCTCTTCCGATC*T |
| DcuSML_FACS_bin-F1_B01_fwd | AATGATACGGCGACCACCGAGATCTACACCCTGGATTGGACACTCTTTCCCTACACGACGCTCTTCCGATC*T |
| DcuSML_FACS_bin-F1_B01_rev | CAAGCAGAAGACGGCATACGAGATCCGCTTCCTAGTGACTGGAGTTCAGACGTGTGCTCTTCCGATC*T |
| DcuSML_FACS_bin-F2_B02_fwd | AATGATACGGCGACCACCGAGATCTACACTCGCGATGTCACACTCTTTCCCTACACGACGCTCTTCCGATC*T |
| DcuSML_FACS_bin-F2_B02_rev | CAAGCAGAAGACGGCATACGAGATAGTAGGATGCGTGACTGGAGTTCAGACGTGTGCTCTTCCGATC*T |
| DcuSML_FACS_bin-F3_B03_fwd | AATGATACGGCGACCACCGAGATCTACACCAATACGACAACACTCTTTCCCTACACGACGCTCTTCCGATC*T |
| DcuSML_FACS_bin-F3_B03_rev | CAAGCAGAAGACGGCATACGAGATACGCTGGACTGTGACTGGAGTTCAGACGTGTGCTCTTCCGATC*T |
| DcuSML_FACS_bin-F4_B04_fwd | AATGATACGGCGACCACCGAGATCTACACCTAGACACTCACACTCTTTCCCTACACGACGCTCTTCCGATC*T |
| DcuSML_FACS_bin-F4_B04_rev | CAAGCAGAAGACGGCATACGAGATATGGCTGCCTGTGACTGGAGTTCAGACGTGTGCTCTTCCGATC*T |
| DcuSML_FACS_bin-F5_B05_fwd | AATGATACGGCGACCACCGAGATCTACACACATGGCCTGACACTCTTTCCCTACACGACGCTCTTCCGATC*T |
| DcuSML_FACS_bin-F5_B05_rev | CAAGCAGAAGACGGCATACGAGATTACGAGTCGCGTGACTGGAGTTCAGACGTGTGCTCTTCCGATC*T |
| DcuSML_FACS_bin-F6_B06_fwd | AATGATACGGCGACCACCGAGATCTACACCAAGATACGTACACTCTTTCCCTACACGACGCTCTTCCGATC*T |
| DcuSML_FACS_bin-F6_B06_rev | CAAGCAGAAGACGGCATACGAGATCCTCAGATACGTGACTGGAGTTCAGACGTGTGCTCTTCCGATC*T |
| DcuSML_FACS_bin-F7_B07_fwd | AATGATACGGCGACCACCGAGATCTACACAATACCACCAACACTCTTTCCCTACACGACGCTCTTCCGATC*T |
| DcuSML_FACS_bin-F7_B07_rev | CAAGCAGAAGACGGCATACGAGATTTAGGTGATGGTGACTGGAGTTCAGACGTGTGCTCTTCCGATC*T |
| DcuSML_FACS_bin-F8_B08_fwd | AATGATACGGCGACCACCGAGATCTACACTCCAGCTCATACACTCTTTCCCTACACGACGCTCTTCCGATC*T |
| DcuSML_FACS_bin-F8_B08_rev | CAAGCAGAAGACGGCATACGAGATGTATATGCCGGTGACTGGAGTTCAGACGTGTGCTCTTCCGATC*T |
| DcuSML_FACS_bin-F9_B09_fwd | AATGATACGGCGACCACCGAGATCTACACCTCTAGCCTAACACTCTTTCCCTACACGACGCTCTTCCGATC*T |
| DcuSML_FACS_bin-F9_B09_rev | CAAGCAGAAGACGGCATACGAGATGACACTCGTGGTGACTGGAGTTCAGACGTGTGCTCTTCCGATC*T |
| DcuSML_FACS_bin-F10_B10_fwd | AATGATACGGCGACCACCGAGATCTACACGTAGAAGCATACACTCTTTCCCTACACGACGCTCTTCCGATC*T |
| DcuSML_FACS_bin-F10_B10_rev | CAAGCAGAAGACGGCATACGAGATATAGTCTCCGGTGACTGGAGTTCAGACGTGTGCTCTTCCGATC*T |
| DcuSML_FACS_bin-F11_B11_fwd | AATGATACGGCGACCACCGAGATCTACACCACCAGCACAACACTCTTTCCCTACACGACGCTCTTCCGATC*T |
| DcuSML_FACS_bin-F11_B11_rev | CAAGCAGAAGACGGCATACGAGATCTAACACCTTGTGACTGGAGTTCAGACGTGTGCTCTTCCGATC*T |
| DcuSML_FACS_bin-F12_B12_fwd | AATGATACGGCGACCACCGAGATCTACACATCTTGCGGCACACTCTTTCCCTACACGACGCTCTTCCGATC*T |
| DcuSML_FACS_bin-F12_B12_rev | CAAGCAGAAGACGGCATACGAGATCTTCCATCGCGTGACTGGAGTTCAGACGTGTGCTCTTCCGATC*T |
| DcuSML_FACS_bin-A1_C01_fwd | AATGATACGGCGACCACCGAGATCTACACTAGAGAGTACACACTCTTTCCCTACACGACGCTCTTCCGATC*T |
| DcuSML_FACS_bin-A1_C01_rev | CAAGCAGAAGACGGCATACGAGATATGCGACCGTGTGACTGGAGTTCAGACGTGTGCTCTTCCGATC*T |
| DcuSML_FACS_bin-A2_C02_fwd | AATGATACGGCGACCACCGAGATCTACACTCTCGGAGTTACACTCTTTCCCTACACGACGCTCTTCCGATC*T |
| DcuSML_FACS_bin-A2_C02_rev | CAAGCAGAAGACGGCATACGAGATCCGTGTGTAAGTGACTGGAGTTCAGACGTGTGCTCTTCCGATC*T |
| DcuSML_FACS_bin-A3_C03_fwd | AATGATACGGCGACCACCGAGATCTACACCCGCGATTAGACACTCTTTCCCTACACGACGCTCTTCCGATC*T |
| DcuSML_FACS_bin-A3_C03_rev | CAAGCAGAAGACGGCATACGAGATTCTAATCTCGGTGACTGGAGTTCAGACGTGTGCTCTTCCGATC*T |
| DcuSML_FACS_bin-A4_C04_fwd | AATGATACGGCGACCACCGAGATCTACACTTGTGTCTGCACACTCTTTCCCTACACGACGCTCTTCCGATC*T |
| DcuSML_FACS_bin-A4_C04_rev | CAAGCAGAAGACGGCATACGAGATTCGATCCTTGGTGACTGGAGTTCAGACGTGTGCTCTTCCGATC*T |
| DcuSML_FACS_bin-A5_C05_fwd | AATGATACGGCGACCACCGAGATCTACACAGTGCCGATGACACTCTTTCCCTACACGACGCTCTTCCGATC*T |
| DcuSML_FACS_bin-A5_C05_rev | CAAGCAGAAGACGGCATACGAGATTGTGTGTTGTGTGACTGGAGTTCAGACGTGTGCTCTTCCGATC*T |
| DcuSML_FACS_bin-A6_C06_fwd | AATGATACGGCGACCACCGAGATCTACACGATCTATTGGACACTCTTTCCCTACACGACGCTCTTCCGATC*T |
| DcuSML_FACS_bin-A6_C06_rev | CAAGCAGAAGACGGCATACGAGATGCGAGCTCTTGTGACTGGAGTTCAGACGTGTGCTCTTCCGATC*T |
| DcuSML_FACS_bin-A7_C07_fwd | AATGATACGGCGACCACCGAGATCTACACCCGATATGTCACACTCTTTCCCTACACGACGCTCTTCCGATC*T |
| DcuSML_FACS_bin-A7_C07_rev | CAAGCAGAAGACGGCATACGAGATACGCCTCTGTGTGACTGGAGTTCAGACGTGTGCTCTTCCGATC*T |
| DcuSML_FACS_bin-A8_C08_fwd | AATGATACGGCGACCACCGAGATCTACACCATATTGGAGACACTCTTTCCCTACACGACGCTCTTCCGATC*T |
| DcuSML_FACS_bin-A8_C08_rev | CAAGCAGAAGACGGCATACGAGATCCGTACAGTTGTGACTGGAGTTCAGACGTGTGCTCTTCCGATC*T |
| DcuSML_FACS_bin-A9_C09_fwd | AATGATACGGCGACCACCGAGATCTACACACACAATGGTACACTCTTTCCCTACACGACGCTCTTCCGATC*T |
| DcuSML_FACS_bin-A9_C09_rev | CAAGCAGAAGACGGCATACGAGATCCTACTTCGCGTGACTGGAGTTCAGACGTGTGCTCTTCCGATC*T |
| DcuSML_FACS_bin-A10_C10_fwd | AATGATACGGCGACCACCGAGATCTACACGATCAAGCCAACACTCTTTCCCTACACGACGCTCTTCCGATC*T |
| DcuSML_FACS_bin-A10_C10_rev | CAAGCAGAAGACGGCATACGAGATGTGTCTTCACGTGACTGGAGTTCAGACGTGTGCTCTTCCGATC*T |
| DcuSML_FACS_bin-A11_C11_fwd | AATGATACGGCGACCACCGAGATCTACACAAGTCCATAGACACTCTTTCCCTACACGACGCTCTTCCGATC*T |
| DcuSML_FACS_bin-A11_C11_rev | CAAGCAGAAGACGGCATACGAGATGACCTAAGTCGTGACTGGAGTTCAGACGTGTGCTCTTCCGATC*T |
| DcuSML_FACS_bin-A12_C12_fwd | AATGATACGGCGACCACCGAGATCTACACCCACGACTTCACACTCTTTCCCTACACGACGCTCTTCCGATC*T |
| DcuSML_FACS_bin-A12_C12_rev | CAAGCAGAAGACGGCATACGAGATTGTCCGTCCTGTGACTGGAGTTCAGACGTGTGCTCTTCCGATC*T |
| DcuSML_Presort1_E01_fwd | AATGATACGGCGACCACCGAGATCTACACCTAACAGCCGACACTCTTTCCCTACACGACGCTCTTCCGATC*T |
| DcuSML_Presort1_E01_rev | CAAGCAGAAGACGGCATACGAGATGACAATCCAAGTGACTGGAGTTCAGACGTGTGCTCTTCCGATC*T |
| DcuSML_Presort2_E03_fwd | AATGATACGGCGACCACCGAGATCTACACACAGACTCATACACTCTTTCCCTACACGACGCTCTTCCGATC*T |
| DcuSML_Presort2_E03_rev | CAAGCAGAAGACGGCATACGAGATGCGATTGCCTGTGACTGGAGTTCAGACGTGTGCTCTTCCGATC*T |
| Mutant_S4A_fwd | AACTCATATGAGACATGCATTGCCCTACC |
| Mutant_S4A_rev | CCTGTCTAGTTGGCCAGGCAGTTAGAA |
| Mutant_M24V_fwd | AACTCATATGAGACATTCATTGCCCTACCG |
| Mutant_M24V_rev | CTCCTGTTTGGTTCCCTTTCTCAATGACA |
| Mutant_Q43R_fwd | AACTCATATGAGACATTCATTGCCCTACCG |
| Mutant_Q43R_rev | GCTTGTGGCGCTATGCAGATGGTT |
| Mutant_V35E_fwd | AACTCATATGAGACATTCATTGCCCTACCG |
| Mutant_V35E_rev | CCTGGTCGAGTGACTAATCGCTACGTT |
| Mutant_V36E_fwd | AACTCATATGAGACATTCATTGCCCTACCG |
| Mutant_V36E_rev | CTCCTCAACTGGGCATTTTCTGCTACATA |
| Mutant_D65E_fwd | AACTCATATGAGACATTCATTGCCCTACCG |
| Mutant_D65E_rev | TTCTCCTACAACTTCCAATAGTGCCTCAAAT |
| Mutant_D65V_fwd | AACTCATATGAGACATTCATTGCCCTACCG |
| Mutant_D65V_rev | CCTTCAATTTGCAACGACTAGCTCGGG |

Table S2. Fluorescence-activated cell sorting bin boundaries in relative fluorescence units (RFU) and molecules of equivalent fluorescein (MEFL).

| **Bin number** | **RFU range** | **MEFL range (No Ligand sort)** | **MEFL range (Fumarate Aspartate sorts)** |
| --- | --- | --- | --- |
| 1 | 100 – 290 | 312 – 836 | 294 – 785 |
| 2 | 290 – 590 | 836 – 1,612 | 785 – 1,511 |
| 3 | 590 – 1,950 | 1,612 – 4,866 | 1,511 – 4,551 |
| 4 | 1,950 – 4,300 | 4,866 – 10,104 | 4,551 – 9,438 |
| 5 | 4,300 – 6,500 | 10,104 – 14,802 | 9,438 – 13,816 |
| 6 | 6,500 – 8,900 | 14,802 – 19,790 | 13,816 – 18,461 |
| 7 | 8,900 – 11,820 | 19,790 – 25722 | 18,641 – 23,982 |
| 8 | 11,820 – 15,820 | 25,722 – 33,674 | 23,982 – 31,379 |
| 9 | 15,820 – 22,450 | 33,674 – 46,533 | 31,379 – 43,334 |
| 10 | 22,450 – 33,000 | 46,533 – 66,429 | 43,334 – 61,819 |
| 11 | 33,000 – 55,000 | 66,429 – 109,504 | 61,819 – 99,021 |
| 12 | 55,000+ | 109,504+ | 99,021+ |

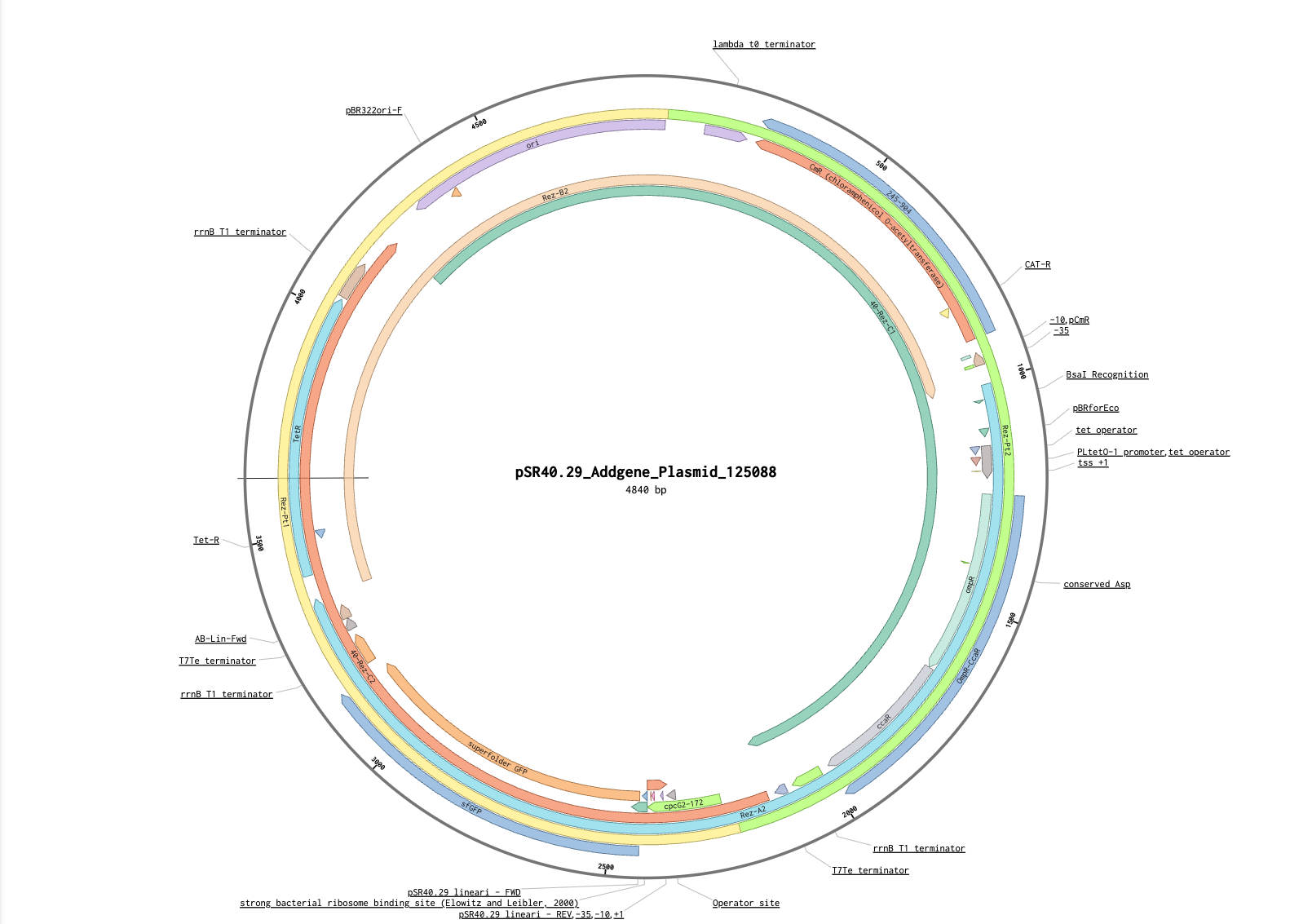

pSR40.29 plasmid map (Addgene plasmid 125088).

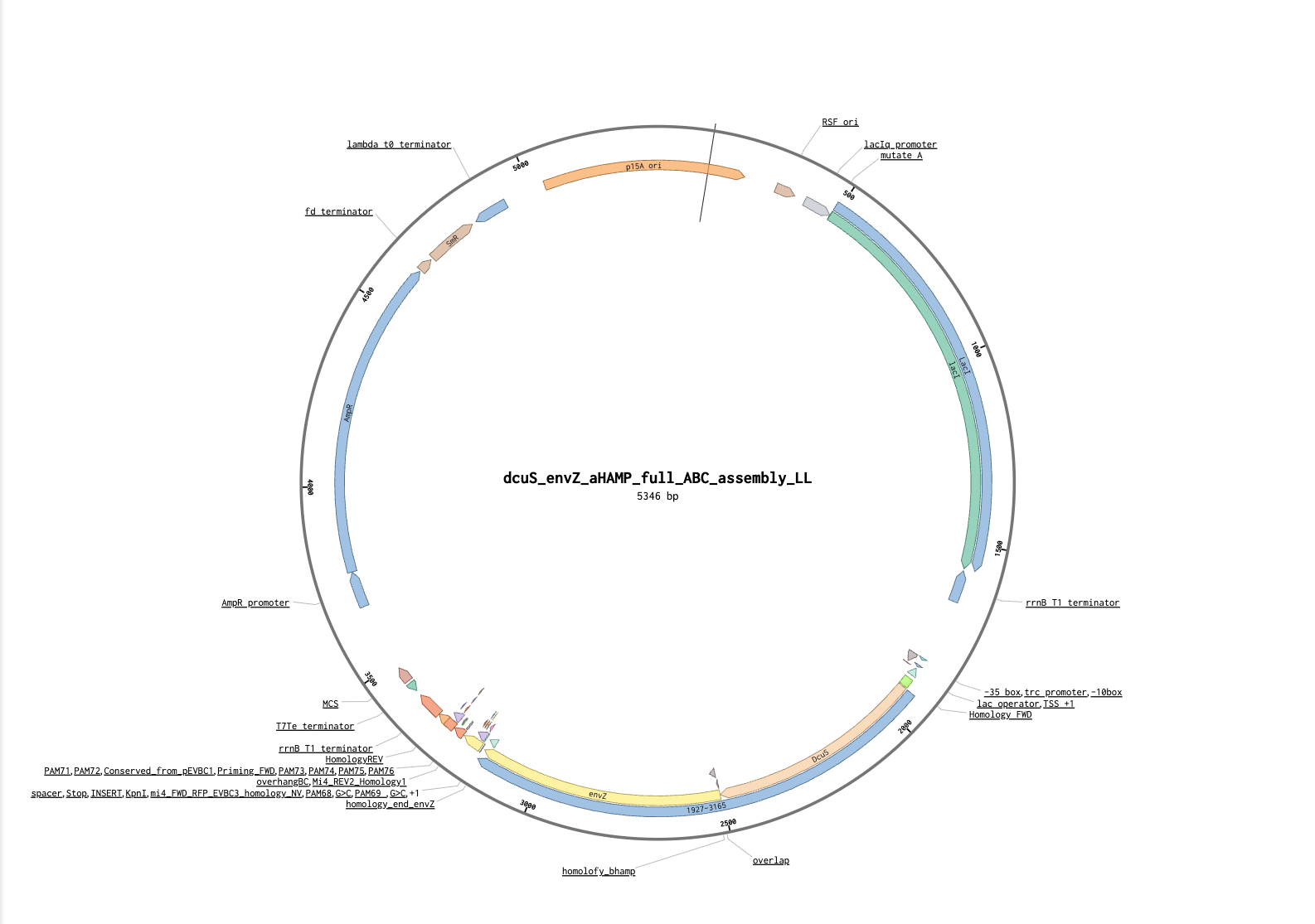

pDcuSEnvZ plasmid map (dcuS_envZ_full_ABC_assembly_LL).

dcuS_envZ_full_ABC_assembly_LL:

acagtccagcttggagcgaactgcctacccggaactgagtgtcaggcgtggaatgagacaaacgcggccataacagcggaatgacaccggtaaaccgaaaggcaggaacaggagagcgcacgagggagccgccagggggaaacgcctggtatctttatagtcctgtcgggtttcgccaccactgatttgagcgtcagatttcgtgatgcttgtcaggggggcggagcctatggaaaaacggctttgccgcggccctctcactttcctgttaagtatcttcctggcatcttccgggaaatctccgccccgttcgtaagccatttccgctcgccgcagtcgaacgaccgagcgtagcgagtcagtgagcgaggaagcggaatatatcccgaagcggcatgcatttacgttgacaccatcgaatggtgcaaaacctttcgcggtatggcatgatagcgcccggaagagagtcaattcagggtggtgaatgtgaaaccagtaacgttatacgatgtcgcagagtatgccggtgtctcttatcagaccgtttcccgcgtggtgaaccaggccagccacgtttctgcgaaaacgcgggaaaaagtggaagcggcgatggcggagctgaattacattccaaaccgcgtggcacaacaactggcgggcaaacagtcgttgctgattggcgttgccacctccagtctggccctgcacgcgccgtcgcaaattgtcgcggcgattaaatctcgcgccgatcaactgggtgccagcgtggtggtgtcgatggtagaacgaagcggcgtcgaagactgtaaagcggcggtgcacaatcttctcgcgcaacgcgtcagtgggctgatcattaactatccgctggatgaccaggatgccattgctgtggaagctgcctgcactaatgttcaggcgttatttcttgatgtctctgaccagacaccaatcaacagtattattttctcccatgaagacggtacgcgactgggcgtggagcatctggtcgcattgggtcaccagcaaatcgcgctgttagcgggcccattaagttctgtctCggcgcgtctgcgtctggctggctggcataaatatctcactcgcaatcaaattcagccgatagcggaacgggaaggcgactggagtgccatgtccggttttcaacaaaccatgcaaatgctgaatgagggcatcgttccaactgcgatgctggttgccaacgatcagatggcgctgggcgcaatgcgcgccattaccgagtccgggctgcgcgttggtgcggatatctcggtagtgggatacgacgatacagaagacagctcatgttatatcccgccgttaaccaccatcaaacaggattttcgcctgctggggcaaaccagcgtggaccgcttgctgcaactctctcagggccaggcggtgaagggcaatcagctgttgcccgtctcactggtgaaaagaaaaaccaccctggcgcccaatacgcaaaccgcctctccccgcgcgttggccgattcattaatgcagctggcacgacaggtttcccgactggaaagcgggcagtgaggcatcaaataaaacgaaaggctcagtcgaaagactgggcctttcgttttatctgttgtttgtcggtgaacgctctcctgagtaggacaaatccgccgccctagacctagggcgttcggctgcggcgagcggtatcagctcactcaaaggcggtaatacgtaaatcactgcataattcgtgtagctcaaggcgcactcccgttctggataatgttttttgcgccgacatcataacggttctggctaatattctgaaatgagctgttgacaattaatcatcggctcgtataatgtgtggaattgtgagcggataacaatttcacacagcactctcctaatttaaatcgcacaactCATatgAGACATTCATTGCCCTACCGCATGTTACGCAAACGTCCGATGAAATTGAGTACCACAGTGATCTTAATGGTCAGTGCGGTACTGTTCTCGGTGCTATTGGTGGTGCATCTGATTTACTTCTCGCAAATCAGTGATATGACGCGAGATGGGCTAGCCAACAAGGCACTGGCAGTGGCGCGTACCCTCGCCGACTCGCCGGAAATCCGTCAGGGCTTGCAGAAAAAACCGCAGGAGAGTGGCATCCAGGCCATCGCGGAAGCCGTACGCAAACGCAACGATCTGCTGTTTATTGTCGTTACCGATATGCAAAGTCTTCGCTACTCGCATCCTGAAGCCCAGCGTATTGGTCAGCCATTTAAAGGTGATGACATCCTTAAAGCGCTGAATGGCGAAGAAAATGTCGCTATCAATCGCGGTTTTCTGGCGCAGGCTTTACGCGTATTTACCCCCATCTACGATGAAAATCATAAACAAATTGGCGTGGTGGCGATCGGCCTTGAGTTAAGCCGTGTGACCCAACAGATCAATGACAGTCGCTGGAGCCTGCAGATGGCGGCTGGTGTTAAGCAACTGGCGgaCgaccgcacgctgctgatggcgggggtaagtcacgacttgcgcacgccgctgacgcgtattcgcctggcgactgagatgatgagcgagcaggatggctatctggcagaatcgatcaataaagatatcgaagagtgcaacgccatcattgagcagtttatcgactacctgcgcaccgggcaggagatgccgatggaaatggcggatcttaatgcagtactcggtgaggtgattgctgccgaaagtggctatgagcgggaaattgaaaccgcgctttaccccggcagcattgaagtgaaaatgcacccgctgtcgatcaaacgcgcggtggcgaatatggtggtcaacgccgcccgttatggcaatggctggatcaaagtcagcagcggaacggagccgaatcgcgcctggttccaggtggaagatgacggtccgggaattgcgccggaacaacgtaagcacctgttccagccgtttgtccgcggcgacagtgcgcgcaccattagcggcacgggattagggctggcaattgtgcagcgtatcgtggataaccataacgggatgctggagcttggcaccagcgagcggggcgggctttccattcgcgcctggctgccagtgccggtaacgcgggcgcagggcacgacaaaagaagggtaaACCTCCCAATAAGGTACCtaaGTGTcGCTGCcGAACagcNNBBDDBBVVHHDDBBVVHNDDNNaggAGAAGAGCgcacGACGTcaCgtCgcaGAATTCcttttcggggaaatgtgcgcggaacccgggtaataataatctagaccaggcatcaaataaaacgaaaggctcagtcgaaagactgggcctttcgttttatctgttgtttgtcggtgaacgctctctactagagtcacactggctcaccttcgggtgggcctttctgcgtttataggtacaggggatcctctagagtcgacctgcaggcatgcaagcttagcaagcgaaccggaattgccagctggggcgccctctggtaaggttgggaagccctgcaaagtaaactggatggctttcttgccgccaaggatctgatggcgcaggggatcaagatctgatcaagagacaggatgaggatcgtttcgtaaattagtagcccgcctaatgagcgggcttttttttaattcccctatttgtttatttttctaaatacattcaaatatgtatccgctcatgagacaataaccctgataaatgcttcaataatattgaaaaaggaagagtatgagcattcagcattttcgtgtggcgctgattccgttttttgcggcgttttgcctgccggtgtttgcgcatccggaaaccctggtgaaagtgaaagatgcggaagatcaactgggtgcgcgcgtgggctatattgaactggatctgaacagcggcaaaattctggaatcttttcgtccggaagaacgttttccgatgatgagcacctttaaagtgctgctgtgcggtgcggttctgagccgtgtggatgcgggccaggaacaactgggccgtcgtattcattatagccagaacgatctggtggaatatagcccggtgaccgaaaaacatctgaccgatggcatgaccgtgcgtgaactgtgcagcgcggcgattaccatgagcgataacaccgcggcgaacctgctgctgacgaccattggcggtccgaaagaactgaccgcgtttctgcataacatgggcgatcatgtgacccgtctggatcgttgggaaccggaactgaacgaagcgattccgaacgatgaacgtgataccaccatgccggcagcaatggcgaccaccctgcgtaaactgctgacgggtgagctgctgaccctggcaagccgccagcaactgattgattggatggaagcggataaagtggcgggtccgctgctgcgtagcgcgctgccggctggctggtttattgcggataaaagcggtgcgggcgaacgtggcagccgtggcattattgcggcgctgggcccggatggtaaaccgagccgtattgtggtgatttataccaccggcagccaggcgacgatggatgaacgtaaccgtcagattgcggaaattggcgcgagcctgattaaacattggtaaaccgatacaattaaaggctccttttggagcctttttttttggacgaccccccagtatcagcccgtcatacttgaagctagacaggcttatcttggacaagaagaagatcgcttggcctcgcgcgcagatcagttggaagaatttgtccactacgtgaaaggcgagatcaccaaggtagtaggcaaataagagctcgcttggactcctgttgatagatccagtaatgacctcagaactccatctggatttgttcagaacgctcggttgccgccgggcgttttttattggtgagaatccaagcactagtaacaacttatatcgtatggggctgacttcaggtgctacatttgaagagataaattgcactgaaatctagtaatattttatctgattaataagatgatcttcttgagatcgttttggtctgcgcgtaatctcttgctctgaaaacgaaaaaaccgccttgcagggcggtttttcgaaggttctctgagctaccaactctttgaaccgaggtaactggcttggaggagcgcagtcaccaaaacttgtcctttcagtttagccttaaccggcgcatgacttcaagactaactcctctaaatcaattaccagtggctgctgccagtggtgcttttgcatgtctttccgggttggactcaagacgatagttaccggataaggcgcagcggtcggactgaacggggggttcgtgcat

Wildtype DcuS/EnvZ chimera sequence:

MRHSLPYRMLRKRPMKLSTTVILMVSAVLFSVLLVVHLIYFSQISDMTRDGLANKALAVARTLADSPEIRQGLQKKPQESGIQAIAEAVRKRNDLLFIVVTDMQSLRYSHPEAQRIGQPFKGDDILKALNGEENVAINRGFLAQALRVFTPIYDENHKQIGVVAIGLELSRVTQQINDSRWSLQMAAGVKQLADDRTLLMAGVSHDLRTPLTRIRLATEMMSEQDGYLAESINKDIEECNAIIEQFIDYLRTGQEMPMEMADLNAVLGEVIAAESGYEREIETALYPGSIEVKMHPLSIKRAVANMVVNAARYGNGWIKVSSGTEPNRAWFQVEDDGPGIAPEQRKHLFQPFVRGDSARTISGTGLGLAIVQRIVDNHNGMLELGTSERGGLSIRAWLPVPVTRAQGTTKEG

DcuS^S4A^/EnvZ amino acid sequence:

MRHALPYRMLRKRPMKLSTTVILMVSAVLFSVLLVVHLIYFSQISDMTRDGLANKALAVARTLADSPEIRQGLQKKPQESGIQAIAEAVRKRNDLLFIVVTDMQSLRYSHPEAQRIGQPFKGDDILKALNGEENVAINRGFLAQALRVFTPIYDENHKQIGVVAIGLELSRVTQQINDSRWSLQMAAGVKQLADDRTLLMAGVSHDLRTPLTRIRLATEMMSEQDGYLAESINKDIEECNAIIEQFIDYLRTGQEMPMEMADLNAVLGEVIAAESGYEREIETALYPGSIEVKMHPLSIKRAVANMVVNAARYGNGWIKVSSGTEPNRAWFQVEDDGPGIAPEQRKHLFQPFVRGDSARTISGTGLGLAIVQRIVDNHNGMLELGTSERGGLSIRAWLPVPVTRAQGTTKEG

DcuS^M24V^/EnvZ amino acid sequence:

MRHSLPYRMLRKRPMKLSTTVILVVSAVLFSVLLVVHLIYFSQISDMTRDGLANKALAVARTLADSPEIRQGLQKKPQESGIQAIAEAVRKRNDLLFIVVTDMQSLRYSHPEAQRIGQPFKGDDILKALNGEENVAINRGFLAQALRVFTPIYDENHKQIGVVAIGLELSRVTQQINDSRWSLQMAAGVKQLADDRTLLMAGVSHDLRTPLTRIRLATEMMSEQDGYLAESINKDIEECNAIIEQFIDYLRTGQEMPMEMADLNAVLGEVIAAESGYEREIETALYPGSIEVKMHPLSIKRAVANMVVNAARYGNGWIKVSSGTEPNRAWFQVEDDGPGIAPEQRKHLFQPFVRGDSARTISGTGLGLAIVQRIVDNHNGMLELGTSERGGLSIRAWLPVPVTRAQGTTKEG

DcuS^Q43R^/EnvZ amino acid sequence:

MRHSLPYRMLRKRPMKLSTTVILMVSAVLFSVLLVVHLIYFSRISDMTRDGLANKALAVARTLADSPEIRQGLQKKPQESGIQAIAEAVRKRNDLLFIVVTDMQSLRYSHPEAQRIGQPFKGDDILKALNGEENVAINRGFLAQALRVFTPIYDENHKQIGVVAIGLELSRVTQQINDSRWSLQMAAGVKQLADDRTLLMAGVSHDLRTPLTRIRLATEMMSEQDGYLAESINKDIEECNAIIEQFIDYLRTGQEMPMEMADLNAVLGEVIAAESGYEREIETALYPGSIEVKMHPLSIKRAVANMVVNAARYGNGWIKVSSGTEPNRAWFQVEDDGPGIAPEQRKHLFQPFVRGDSARTISGTGLGLAIVQRIVDNHNGMLELGTSERGGLSIRAWLPVPVTRAQGTTKEG

DcuS^V35E^/EnvZ amino acid sequence:

MRHSLPYRMLRKRPMKLSTTVILMVSAVLFSVLLEVHLIYFSQISDMTRDGLANKALAVARTLADSPEIRQGLQKKPQESGIQAIAEAVRKRNDLLFIVVTDMQSLRYSHPEAQRIGQPFKGDDILKALNGEENVAINRGFLAQALRVFTPIYDENHKQIGVVAIGLELSRVTQQINDSRWSLQMAAGVKQLADDRTLLMAGVSHDLRTPLTRIRLATEMMSEQDGYLAESINKDIEECNAIIEQFIDYLRTGQEMPMEMADLNAVLGEVIAAESGYEREIETALYPGSIEVKMHPLSIKRAVANMVVNAARYGNGWIKVSSGTEPNRAWFQVEDDGPGIAPEQRKHLFQPFVRGDSARTISGTGLGLAIVQRIVDNHNGMLELGTSERGGLSIRAWLPVPVTRAQGTTKEG

DcuS^V36E^/EnvZ amino acid sequence:

MRHSLPYRMLRKRPMKLSTTVILMVSAVLFSVLLVEHLIYFSQISDMTRDGLANKALAVARTLADSPEIRQGLQKKPQESGIQAIAEAVRKRNDLLFIVVTDMQSLRYSHPEAQRIGQPFKGDDILKALNGEENVAINRGFLAQALRVFTPIYDENHKQIGVVAIGLELSRVTQQINDSRWSLQMAAGVKQLADDRTLLMAGVSHDLRTPLTRIRLATEMMSEQDGYLAESINKDIEECNAIIEQFIDYLRTGQEMPMEMADLNAVLGEVIAAESGYEREIETALYPGSIEVKMHPLSIKRAVANMVVNAARYGNGWIKVSSGTEPNRAWFQVEDDGPGIAPEQRKHLFQPFVRGDSARTISGTGLGLAIVQRIVDNHNGMLELGTSERGGLSIRAWLPVPVTRAQGTTKEG

DcuS^D65E^/EnvZ amino acid sequence:

MRHSLPYRMLRKRPMKLSTTVILMVSAVLFSVLLVVHLIYFSQISDMTRDGLANKALAVARTLAESPEIRQGLQKKPQESGIQAIAEAVRKRNDLLFIVVTDMQSLRYSHPEAQRIGQPFKGDDILKALNGEENVAINRGFLAQALRVFTPIYDENHKQIGVVAIGLELSRVTQQINDSRWSLQMAAGVKQLADDRTLLMAGVSHDLRTPLTRIRLATEMMSEQDGYLAESINKDIEECNAIIEQFIDYLRTGQEMPMEMADLNAVLGEVIAAESGYEREIETALYPGSIEVKMHPLSIKRAVANMVVNAARYGNGWIKVSSGTEPNRAWFQVEDDGPGIAPEQRKHLFQPFVRGDSARTISGTGLGLAIVQRIVDNHNGMLELGTSERGGLSIRAWLPVPVTRAQGTTKEG

DcuS^D65V^/EnvZ amino acid sequence:

MRHSLPYRMLRKRPMKLSTTVILMVSAVLFSVLLVVHLIYFSQISDMTRDGLANKALAVARTLAVSPEIRQGLQKKPQESGIQAIAEAVRKRNDLLFIVVTDMQSLRYSHPEAQRIGQPFKGDDILKALNGEENVAINRGFLAQALRVFTPIYDENHKQIGVVAIGLELSRVTQQINDSRWSLQMAAGVKQLADDRTLLMAGVSHDLRTPLTRIRLATEMMSEQDGYLAESINKDIEECNAIIEQFIDYLRTGQEMPMEMADLNAVLGEVIAAESGYEREIETALYPGSIEVKMHPLSIKRAVANMVVNAARYGNGWIKVSSGTEPNRAWFQVEDDGPGIAPEQRKHLFQPFVRGDSARTISGTGLGLAIVQRIVDNHNGMLELGTSERGGLSIRAWLPVPVTRAQGTTKEG
